# Auto-inhibition of Cnn binding to γ-TuRCs prevents ectopic microtubule nucleation and cell division defects

**DOI:** 10.1101/2020.10.05.326587

**Authors:** Corinne A. Tovey, Chisato Tsuji, Alice Egerton, Fred Bernard, Antoine Guichet, Marc de la Roche, Paul T. Conduit

## Abstract

γ-tubulin ring complexes (γ-TuRCs) nucleate microtubules. They are recruited to centrosomes in dividing cells via binding to N-terminal CM1 domains within γ-TuRC-tethering proteins, including *Drosophila* Cnn. Binding promotes microtubule nucleation and is restricted to centrosomes in dividing cells, but the mechanism regulating binding remains unknown. Here we identify an extreme N-terminal “CM1 auto-inhibition” (CAI) domain found specifically within the centrosomal isoform of Cnn (Cnn-C) that inhibits γ-TuRC binding. Robust binding occurs after removal of the CAI domain or with the addition of phospho-mimetic mutations, suggesting that phosphorylation helps relieve inhibition. We show that regulation of Cnn binding to γ-TuRCs is isoform-specific and that mis-regulation of binding can result in ectopic cytosolic microtubules and major defects during cell division. We also find that human CDK5RAP2 is auto-inhibited from binding γ-TuRCs, suggesting conservation across species. Overall, our results shed light on how and why CM1 domain binding to γ-TuRCs is regulated.

**Summary (for the online JCB table of contents and alerts):** We show that auto-inhibition regulates the binding between microtubule nucleating complexes and proteins that tether them to sites of microtubule nucleation. Failure to properly regulate this binding can lead to ectopic cytosolic microtubule nucleation and major defects during cell division.

## Introduction

Microtubules are polarised polymers of tubulin that are organised into specialised arrays crucial for cell function, such as the mitotic spindle. Correct array assembly relies in part on the spatiotemporal regulation of microtubule formation, and this is achieved by restricting microtubule formation and organisation to specific microtubule organising centres (MTOCs), such as the centrosome during mitosis (Tillery et al., 2018; Sanchez and Feldman, 2016; Petry and Vale, 2015).

The common link between most MTOCs is the presence of multi-protein γ-tubulin ring complexes (γ-TuRCs), which template and catalyse the kinetically unfavourable process of microtubule nucleation (Kollman et al., 2011; Teixidó-Travesa et al., 2012; Lin et al., 2014a; Tovey and Conduit, 2018; Farache et al., 2018). γ-TuRCs are recruited to MTOCs by γ-TuRC-tethering proteins that directly link γ-TuRCs to the MTOC. γ-TuRCs contain 14 γ-tubulin molecules held in a single-turn helical conformation by laterally associating γ-tubulin complex proteins (GCPs) that bind directly to *α*/*β*-tubulin dimers to promote new microtubule assembly. γ-TuRCs have a low activity within the cytosol but are thought to be “activated” after recruitment to MTOCs. In this model, the controlled recruitment and activation of γ-TuRCs enables the spatiotemporal control of microtubule nucleation and array formation. Consistent with this model, recent structural studies have shown that γ-TuRCs purified from the cytosol of HeLa cells and *Xenopus* eggs are in a semi-open conformation, in which the γ-tubulin molecules do not perfectly match the geometry of a 13 protofilament microtubule (Consolati et al., 2020; Liu et al., 2019; Wieczorek et al., 2019). A conformational change into a fully closed ring that matches the geometry of a microtubule is expected to increase the nucleation capacity of the γ-TuRC. This agrees with studies in budding yeast, where there are conformational differences between γ-TuRC-like structures formed *in vitro* and γ-TuRCs bound to microtubules *in vivo*, and where artificial “closure” of γ-TuRCs increases microtubule nucleation capacity (Kollman et al., 2015).

How activation via an open-to-closed conformation change occurs is currently unclear, but various factors have been reported to increase nucleation capacity. γ-TuRCs purified from *Xenopus* egg extract nucleate much more efficiently after the addition of the TOG domain protein XMAP215 (Thawani et al., 2020). TOG domain family members mediate *α*/*β*-tubulin addition via their TOG domains (Nithianantham et al., 2018), bind directly to γ-tubulin, and function in microtubule nucleation *in vitro* and *in vivo* (Wieczorek et al., 2015; Roostalu et al., 2015; Thawani et al., 2018; Flor-Parra et al., 2018; Gunzelmann et al., 2018). Single molecule experiments combined with modelling suggest that XMAP215 indirectly promotes the open-to-closed conformation change of purified γ-TuRCs by increasing the chance of protofilament formation, as lateral contacts between protofilaments should promote γ-TuRC closure (Thawani et al., 2020). While this is an attractive model, evidence suggests that activation can occur in different ways and may be context specific. Phosphorylation of γ-TuRCs by Aurora A around mitotic chromatin increases γ-TuRC activity (Pinyol et al., 2013; Scrofani et al., 2015), as does addition of NME7 kinase *in vitro* (Liu et al., 2014). γ-TuRC activity is also increased after binding of the Augmin complex (Tariq et al., 2020), which tethers γ-TuRCs to other microtubules.

Another well-documented potential γ-TuRC activator is the Centrosomin Motif 1 (CM1) domain, which is conserved in γ-TuRC-tethering proteins across Eukaryotes (Sawin et al., 2004; Zhang and Megraw, 2007; Lin et al., 2014b). Addition of protein fragments containing the CM1 domain increase the nucleation capacity of γ-TuRCs purified from human cells (Choi et al., 2010; Muroyama et al., 2016), although the degree of this activity change was much lower or absent when using γ-TuRCs purified from *Xenopus* eggs (Liu et al., 2019; Thawani et al., 2020). Expression of CM1 domain fragments within human cells leads to the ectopic nucleation of microtubules throughout the cytosol, and this is dependent on CM1 binding to γ-TuRCs (Choi et al., 2010; Hanafusa et al., 2015; Cota et al., 2017). In fission yeast, expression of CM1 domain fragments also results in cytosolic microtubule nucleation (Lynch et al., 2014), and in *Xenopus* addition of CM1-domain fragments increases microtubule aster formation within egg extracts supplemented with activated Ran (Liu et al., 2019). In budding yeast, CM1 domain binding appears to move γ-tubulin molecules into a better position for nucleation (Brilot et al., 2019). While large global structural changes were not observed in mammalian γ-TuRCs bound by the CM1 domain (Liu et al., 2019; Wieczorek et al., 2019), local structural changes can be observed, suggesting that more global changes could in theory occur with a higher stoichiometry of binding (Brilot et al., 2019).

Given that CM1-domain binding leads to microtubule nucleation, binding is likely spatiotemporally controlled, particularly during cell division. This idea is consistent with results from numerous mass spectrometry experiments showing that γ-TuRCs do not readily associate with CM1-domain proteins within the cytosol (Oegema et al., 1999; Choi et al., 2010; Hutchins et al., 2010; Teixidó-Travesa et al., 2012; Thawani et al., 2018; Liu et al., 2019; Wieczorek et al., 2019; Consolati et al., 2020). Binding of the human and *C. elegans* CM1 domain proteins, CDK5RAP2 and SPD-5, to γ-TuRCs involves phosphorylation (Hanafusa et al., 2015; Ohta et al., 2021), which can be a means to spatiotemporally control binding. Nevertheless, whether phosphorylation directly promotes binding to γ-TuRCs or regulates binding in a different way remains unclear.

*Drosophila* Centrosomin (Cnn) is the only reported CM1-domain protein in *Drosophila* but is a multi-isoform gene with all isoforms containing the CM1 domain (Eisman et al., 2009). The centrosomal isoform (Cnn-C) has a dual role, both in recruiting γ-TuRCs to centrosomes (Zhang and Megraw, 2007; Conduit et al., 2014b) and in forming a centrosome-scaffold that supports mitotic pericentriolar material assembly (Conduit et al., 2014a; Feng et al., 2017). Phosphorylation of a central Phospho-regulated multimerisation (PReM) domain specifically at centrosomes drives the oligomerisation of Cnn-C molecules into a scaffold-like structure that helps recruit other centrosomal proteins (Conduit et al., 2014a; Feng et al., 2017). Testes-specific Cnn-T isoforms have mitochondrial localisation domains instead of the PReM and CM2 domains and recruits γ-TuRCs to mitochondria in sperm cells (Chen et al., 2017). Cnn-C and Cnn-T also vary in their extreme N-terminal regions, upstream of the CM1 domain, with Cnn-C containing a longer sequence.

Here, we show that the longer extreme N-terminal region of Cnn-C inhibits binding to γ-TuRCs and therefore name this region the CM1 auto-inhibition (CAI) domain. Removal of the CAI domain leads to robust binding, similar to the robust binding observed for the N-terminal region of Cnn-T. We identify two putative phosphorylation sites, one in the CAI domain (T^27^) and one downstream of the CM1 domain (S^186^), that promote binding to γ-TuRCs when phospho-mimicked, suggesting that phosphorylation relieves CAI domain auto-inhibition. We show that auto-inhibition is important, as expressing a form of Cnn that binds to cytosolic γ-TuRCs leads to cytosolic microtubule nucleation and major defects during cell division. We further show that human CDK5RAP2 is inhibited from binding γ-TuRCs in the cytosol by a region downstream of the CM1 domain, showing that auto-inhibition of binding is conserved feature of CM1 domain proteins.

## Results

### The extreme N-terminal region of Cnn-C is inhibitory for γ-TuRC binding

We previously published evidence that different isoforms of Cnn bind γ-TuRCs with different affinities (Tovey et al., 2018). We found that bacterially purified MBP-tagged N-terminal fragments of Cnn-T (MBP-Cnn-T-N) could immunoprecipitate cytosolic γ-tubulin with a much higher affinity than the equivalent fragments of Cnn-C (MBP-Cnn-C-N). Both isoforms share a short sequence just proximal to the CM1 domain (residues 78-97 in Cnn-C), but differ in their extreme N-terminal region, which is 77 and 19 residues in Cnn-C and Cnn-T, respectively (Figure 1A). We had hypothesised that the larger extreme N-terminal region of Cnn-C may auto-inhibit the CM1 domain, restricting its ability to bind γ-TuRCs. To address this directly, and to confirm the *in vitro* results, we developed an *in vivo* assay where γ-TuRC recruitment to different types of Cnn “scaffolds” formed within eggs could be monitored. To form scaffolds within eggs we injected *in vitro*-generated mRNA encoding Cnn-C with phospho-mimetic mutations within the PReM domain (Cnn-C-PReM^m^) (Figure 1B). The mRNA is translated into protein within the egg and the phospho-mimetic mutations cause the Cnn molecules to oligomerise into centrosome-like scaffolds throughout the cytosol (Conduit et al., 2014a) (Figure 1C-F; Figure S1). To investigate how binding between Cnn and γ-TuRCs is regulated, we modified the N-terminal region of Cnn-C-PReM^m^ (Figure 1B) and measured how efficiently fluorescently-tagged γ-TuRC proteins could be recruited to the scaffolds.

**Figure 1.**
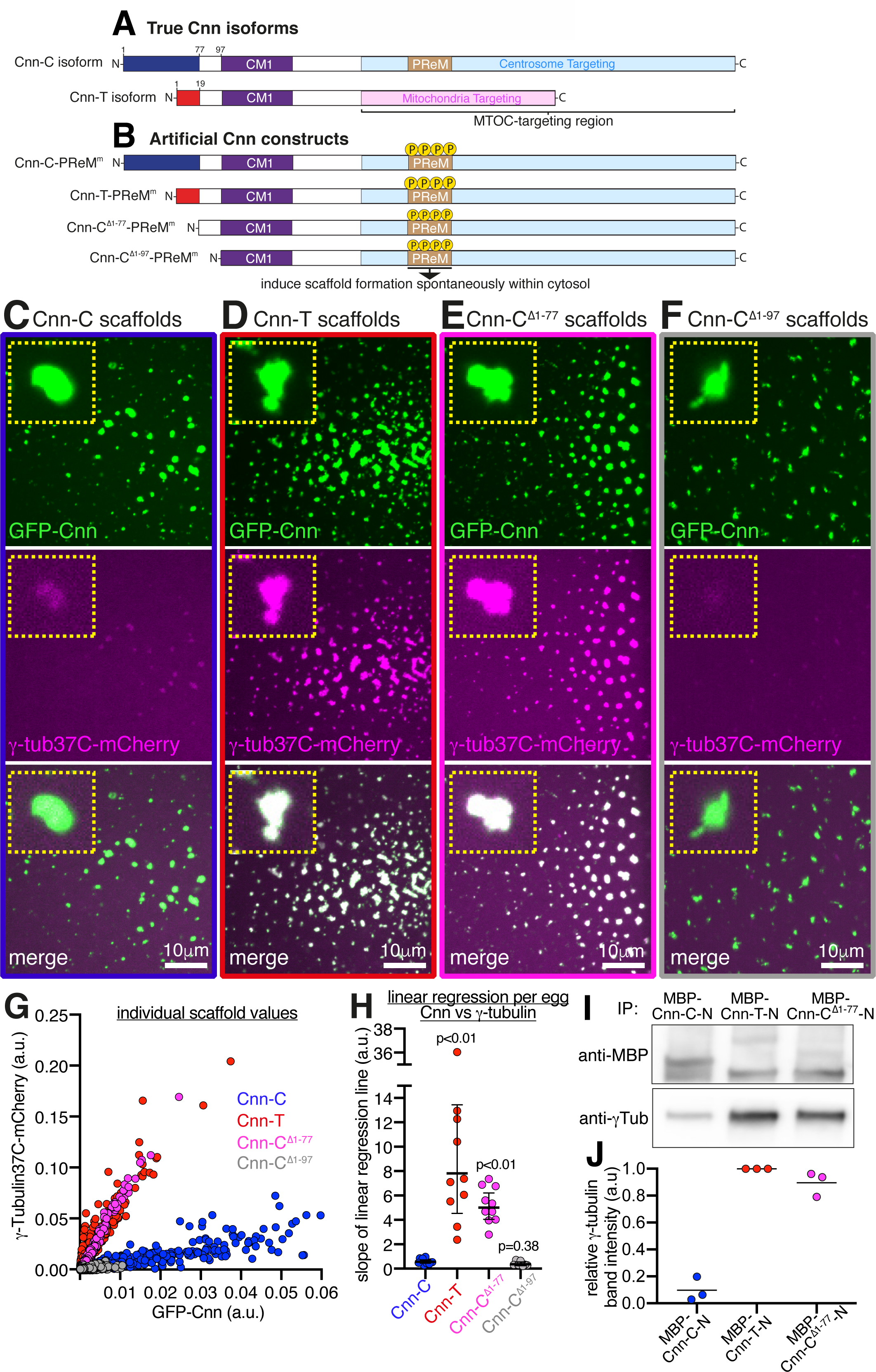
The extreme N-terminal region of Cnn-C inhibits binding to γ-tubulin complexes. (**A**) Diagram of the centrosomal Cnn (Cnn-C) and testes-specific Cnn (Cnn-T) isoforms that exist *in vivo*. **(B)** Diagram of artificial Cnn proteins with differing N-terminal regions used to form Cnn scaffolds (induced by phospho-mimetic mutations in the PReM domain (beige)) via mRNA injection into unfertilised eggs. **(C-F)** Fluorescence images of unfertilised eggs expressing γ-tubulin37C-mCherry that were injected with mRNA encoding different types of artificial Cnn protein, as indicated. Insets show representative examples of individual scaffolds. **(G)** Graph showing fluorescence intensity measurements (in arbitrary units) of γ-tubulin37C-mCherry and GFP-Cnn at Cnn-C (n= 1498 scaffolds; 12 eggs), Cnn-T (n= 1400 scaffolds; 10 eggs), Cnn-C^Δ1-77^ (n= 2168 scaffolds; 10 eggs), or Cnn-C^Δ1-97^ (n= 400 scaffolds; 7 eggs) scaffolds. Each dot represents a single scaffold. **(H)** Graph shows slope values of linear regression lines calculated for scaffolds of different types. Each slope value represents an individual egg that contained multiple scaffolds. The geometric mean and 95% CIs are indicated. P values are from comparisons to the Cnn-C mean using a one-way ANOVA of log_10_ transformed data. (**I**,**J**) Western blot of a co-IP experiment (I) and quantification of γ-tubulin bands (J) showing the efficiency with which different MBP-tagged N-terminal fragments of Cnn, as indicated, co-IP γ-tubulin from embryo extracts. γ-tubulin band intensities were normalised within each of 3 experimental repeats to the γ-tubulin band in the respective MBP-Cnn-T-N IP.

We first compared the recruitment of endogenously tagged γ-tubulin37C-mCherry to GFP-tagged scaffolds formed from unmodified Cnn-C-PReM^m^ with recruitment to scaffolds where the extreme N-terminal region (dark blue in Figure 1A,B) was either exchanged with the extreme N-terminal region of Cnn-T (red in Figure 1A,B) (Cnn-T-PReM^m^) or was removed (Cnn-C^Δ1-77^-PReM^m^). We also tested scaffolds in which all N-terminal amino acids up until the start of the CM1 domain were removed (Cnn-C^Δ1-97^-PReM^m^). For simplicity we refer to these as Cnn-C, Cnn-T, Cnn-C^Δ1-77^, and Cnn-C^Δ1-97^ scaffolds, respectively, regardless of the fluorescent tag used. Initial observations suggested that γ-tubulin associated much more readily with Cnn-T and Cnn-C^Δ1-77^ scaffolds than with Cnn-C or Cnn-C^Δ1-97^ scaffolds (Figure 1C-F). This was clear after plotting the GFP (Cnn) and mCherry (γ-tubulin) fluorescence values for individual scaffolds from multiple embryos per condition (Figure 1G). To quantify γ-tubulin recruitment we performed linear regression for each egg separately and plotted the slope of these lines (S values, in arbitrary units). The mean S value provides an estimate for the relative binding affinity between the different forms of Cnn and γ-tubulin complexes (Figure 1H). The mean S values for Cnn-T scaffolds (7.81) and Cnn-C^Δ1-77^ scaffolds (5.01) were ∼13-fold and 9-fold higher, respectively, than the mean S value for Cnn-C scaffolds (0.57). Consistent with this, MBP-tagged N-terminal fragments of Cnn-T (MBP-Cnn-T-N) and Cnn-C^Δ1-77^ (MBP-Cnn-C-N^Δ1-77^) both co-immunoprecipitated more γ-tubulin from embryo extracts than N-terminal fragments of Cnn-C (MBP-Cnn-C-N) (Figure 1I,J). Thus, the extreme N-terminal region of Cnn-C (blue in Figure 1A,B) is inhibitory for binding to γ-tubulin complexes.

The ability of Cnn-C^Δ1-77^ to bind γ-tubulin complexes appeared to be dependent on the amino acids just upstream of the CM1 domain (aa78-97), which are shared with Cnn-T (Figure 1A), as the mean S value for Cnn-C^Δ1-97^ scaffolds (0.36) was not significantly different from the mean S value for Cnn-C scaffolds (0.57) (Figure 1H). This is consistent with recent observations in *S. cerevisiae*, showing that the equivalent amino acids within the CM1 domain protein SPC110 make close contacts with SPC98^GCP3^ (Brilot et al., 2019).

Cnn-T and Cnn-C^Δ1-77^ scaffolds also recruited the γ-TuRC-specific component Grip75^GCP4^ better than Cnn-C scaffolds (Figure 2A-E). Similar to the recruitment of γ-tubulin, the mean S values for Cnn-T (3.8) and Cnn-C^Δ1-77^ (3.1) scaffolds were 10.3-fold and 8.4-fold higher, respectively, than the S value for Cnn-C (0.37) scaffolds (Figure 2E). Moreover, a combination of western blotting and mass spectrometry showed that bacterially purified MBP-Cnn-T-N fragments could co-immunoprecipitate numerous other γ-TuRC components (Figure S2).

**Figure 2.**
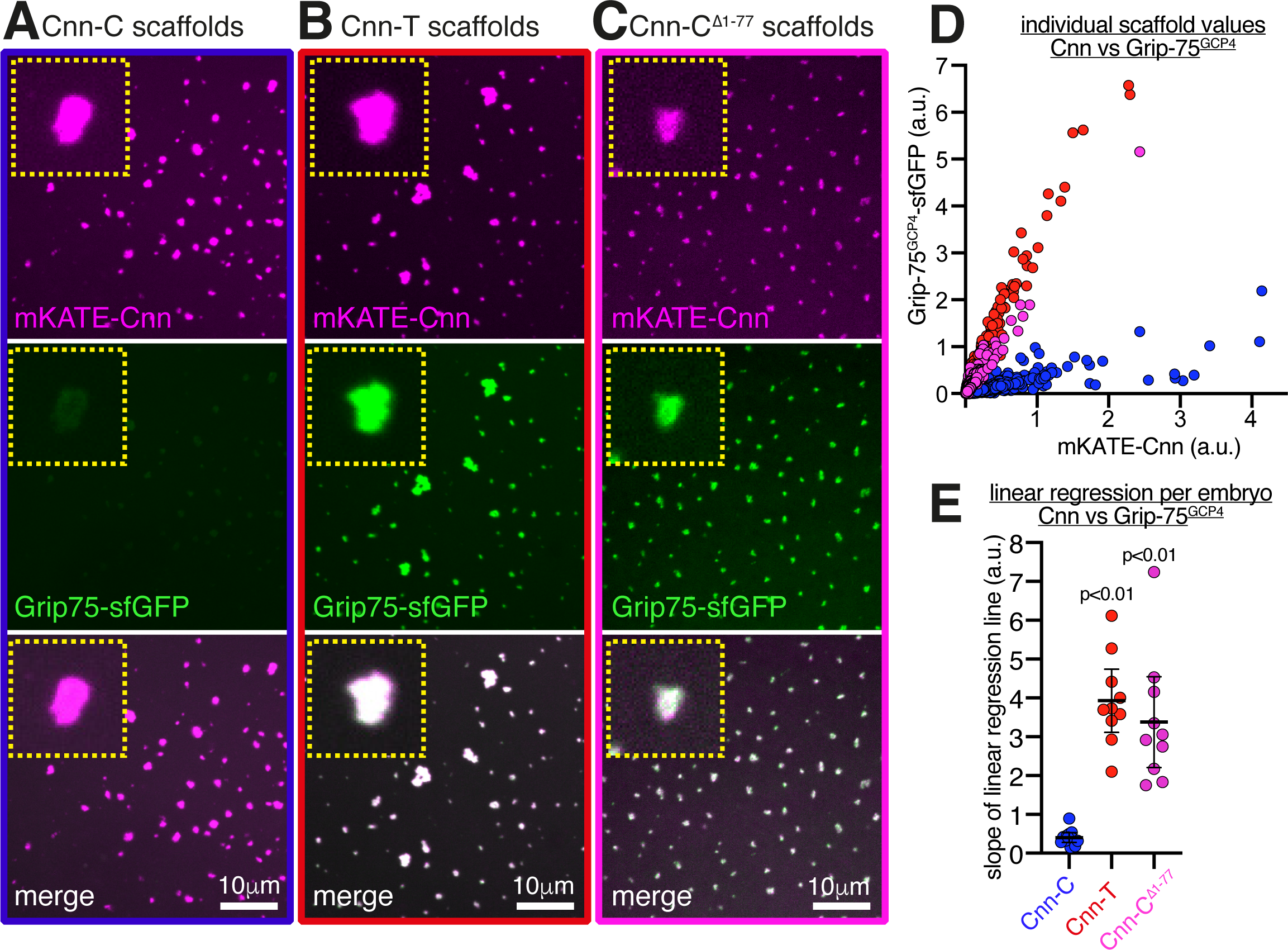
The γ-TuRC-specific protein Grip75^GCP4^ is recruited strongly to Cnn-T and Cnn-C^Δ1-77^ scaffolds. Fluorescence images **(A-C)** show mKATE-Cnn scaffolds of different types, as indicated, within eggs expressing endogenously-tagged Grip75^GCP4^-sfGFP. Insets in show representative examples of individual scaffolds. (**D**) Graph showing fluorescence intensity measurements (in arbitrary units) of Grip75^GCP4^-sfGFP and mKATE-Cnn at Cnn-C (n= 1920 scaffolds; 12 eggs), Cnn-T (n= 1650 scaffolds; 10 eggs) or Cnn-C^Δ1-77^ (n= 2599 scaffolds; 10 eggs). Each dot represents a single scaffold. **(E)** Graph shows slope values of linear regression lines calculated for scaffolds of different types. Each slope value represents an individual egg that contained multiple scaffolds. The mean and 95% CIs are indicated. P values are from comparisons to the Cnn-C mean using a one-way ANOVA.

The data collectively show that the N-terminus of Cnn-T binds robustly to γ-TuRCs, while the extreme N-terminal region of Cnn-C (aa1-77) inhibits binding to γ-TuRCs. We therefore name this region the “CM1 auto-inhibition” (CAI) domain.

### γ-TuRCs recruited by Cnn scaffolds appear to be functional and can generate dynamic microtubules

We next compared the ability of different scaffold types to organise microtubules. We imaged GFP-tagged Cnn-C (low γ-TuRC binding), Cnn-T, or Cnn-C^Δ1-77^ (high γ-TuRC binding) scaffolds within eggs expressing the microtubule binding protein Jupiter-mCherry (Figure 3A-C) and performed a blind analysis to categorise eggs into those containing scaffolds that organised strong, weak, or no microtubule asters (Figure 3D). We also included a “tubulin overlay” category, where the Jupiter-mCherry signal did not extend beyond the GFP scaffold signal. The results show that Cnn-T and Cnn-C^Δ1-77^ scaffolds were much more likely to organise microtubule asters than Cnn-C scaffolds (Figure 3D). This correlates with the increased recruitment of γ-TuRCs to Cnn-T and Cnn-C^Δ1-77^ scaffolds (Figure 1H), suggesting that these γ-TuRCs are able to nucleate microtubules. While it is possible that some microtubules could have been generated independently of γ-TuRCs, a process that occurs by tubulin concentration at *C. elegans* SPD-5 condensates formed *in vitro* (Woodruff et al., 2017), the increased microtubule organising capacity at Cnn-T and Cnn-C^Δ1-77^ scaffolds (high γ-TuRC recruitment) compared to at Cnn-C scaffolds (low γ-TuRC recruitment) suggests that γ-TuRC-mediated microtubule nucleation/organisation is the predominant factor at these Cnn scaffolds.

**Figure 3.**
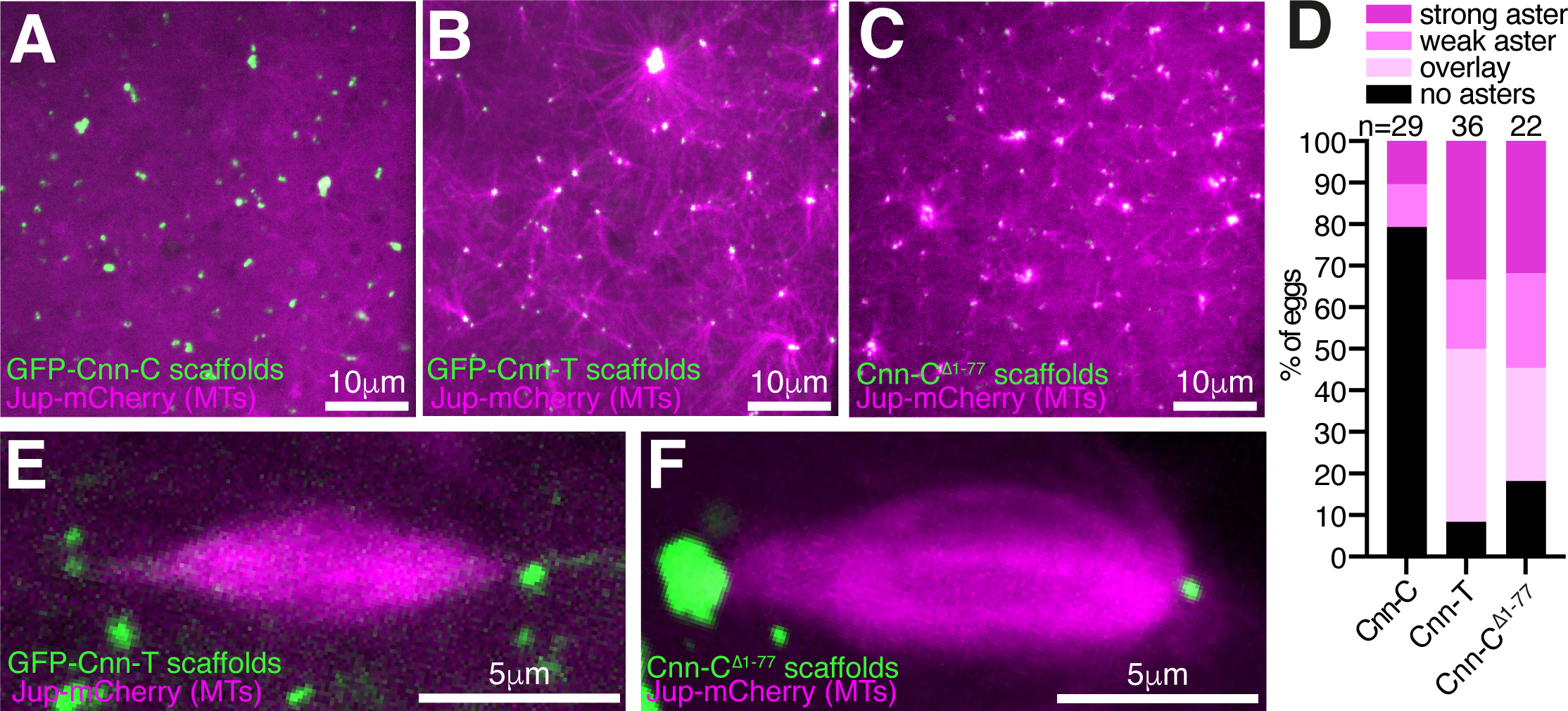
Cnn-T and Cnn-C^Δ1-77^ scaffolds organise microtubules more robustly than Cnn-C scaffolds. **(A-C)** Fluorescence images of Cnn-C scaffolds (A), Cnn-T scaffolds (B), or Cnn-C^Δ1-77^ scaffolds **(C)** within eggs expressing the microtubule marker Jupiter-mCherry. (**D**) Bar-graph showing results of a blind categorisation of eggs containing the different scaffold types based on the ability of the scaffolds within each egg to organise microtubule asters (numbers of eggs analysed indicated above). **(E,F)** Fluorescence images showing that adjacent Cnn-T (E) or Cnn-C^Δ1-77^ (**F**) scaffolds can organise spindle-like structures.

Filming Cnn-T scaffolds through time revealed that these scaffolds could merge and could also be quite mobile, especially those that had microtubules emanating from just one side (Video 1). We could also observe events where spindle-like structures formed between adjacent Cnn-T or Cnn-C^Δ1-77^ scaffolds (Figure 3D; Video 2 and 3), suggesting that the microtubules are dynamic and can be regulated by motor proteins. Giant Cnn-T scaffolds that rotated dragged their attached microtubules through the cytosol, indicating that the microtubules were robustly anchored to the scaffolds (Video 4). In summary, Cnn-T scaffolds can recruit γ-TuRCs that are capable of nucleating and anchoring microtubules.

### Phospho-mimetic mutations help relieve CAI domain mediated auto-inhibition

How could CAI domain mediated auto-inhibition be relieved to allow efficient binding to γ-TuRCs at MTOCs? Studies in human cells, *C. elegans*, and *S. cerevisiae* have shown that binding of CM1 domain proteins to γ-TuSCs or γ-TuRCs is promoted by phosphorylation close to the CM1 domain (Hanafusa et al., 2015; Ohta et al., 2021; Lin et al., 2014b) (Figure S3B). Moreover, Cnn-C binds γ-TuRCs and is phosphorylated only at centrosomes (Zhang and Megraw, 2007; Conduit et al., 2014b; a), suggesting a possible link between binding and phosphorylation.

In an attempt to find phosphorylation sites that may relive CAI domain inhibition, we aligned amino acids 1 to ∼255 of Cnn-C homologues from various *Drosophila* species. We identified three putative phosphorylation “patches” (P1, P2, P3) based on a high concentration of conserved serine and threonine residues (Figure 4A; Figure S3). P1 represented the only region within the CAI domain with predicted secondary structure, corresponding to an *α*-helix (Figure S3). We compared the amount of γ-tubulin that co-IP’d with purified MBP-tagged N-terminal fragments of Cnn-C containing phospho-mimetic mutations (S>D or T>E) in all serine and threonine residues within either P1 (MBP-Cnn-C-N^P1^), P2 (MBP-Cnn-C-N^P2^), P3 (MBP-Cnn-C-N^P3^), or in all three patches (MBP-Cnn-C-N^P1-3^). The original MBP-Cnn-C-N (low binding) and MBP-Cnn-T-N (high binding) fragments were included as negative and positive controls, respectively. Of these phospho-mimetic fragments, MBP-Cnn-C-N^P1^ co-IP’d γ-tubulin most efficiently, although not as efficiently as MBP-Cnn-T-N (Figure 4B,D). We therefore generated phospho-mimetic fragments where either the proximal (S^21^, S^22^, T^27^) or distal (T^31^, T^33^, S^34^) three residues within P1 were mimicked (MBP-Cnn-C-N^P1a^ or MBP-Cnn-C-N^P1b^, respectively). We also phospho-mimicked T^27^ alone (MBP-Cnn-C-N^T27^), because T^27^ is a putative Polo/Plk1 site and because a previous study reported centrosome defects when this site was mutated *in vivo* (Eisman et al., 2015). MBP-Cnn-C-N^P1a^ and MBP-Cnn-C-N^T27^, but not MBP-Cnn-C-N^P1b^, co-IP’d more γ-tubulin than MBP-Cnn-C-N, although again not as much as MBP-Cnn-T-N (Figure 4C,D). In the scaffold assay, phospho-mimicking T^27^ also had a positive effect that was not as strong as that seen with Cnn-T or Cnn-C^Δ1-77^ scaffolds. The mean S value for Cnn-C^T27^ scaffolds (1.35) was ∼2.4-fold higher than for Cnn-C scaffolds (0.57) but still lower than the S values for Cnn-T or Cnn-C^Δ1-77^ scaffolds (Figure 4G; note that S values for Cnn-C^T27E^ scaffolds, and subsequent scaffolds analysed below, were compared to the S values for Cnn-C, Cnn-T and Cnn-C^Δ1-77^ scaffolds from Figure 1H). Together, this suggested that while phosphorylation of T^27^ may be involved in relieving CAI domain auto-inhibition (or in directly increasing the binding affinity between Cnn-C and γ-TuRCs), it is not sufficient for robust γ-TuRC binding.

**Figure 4.**
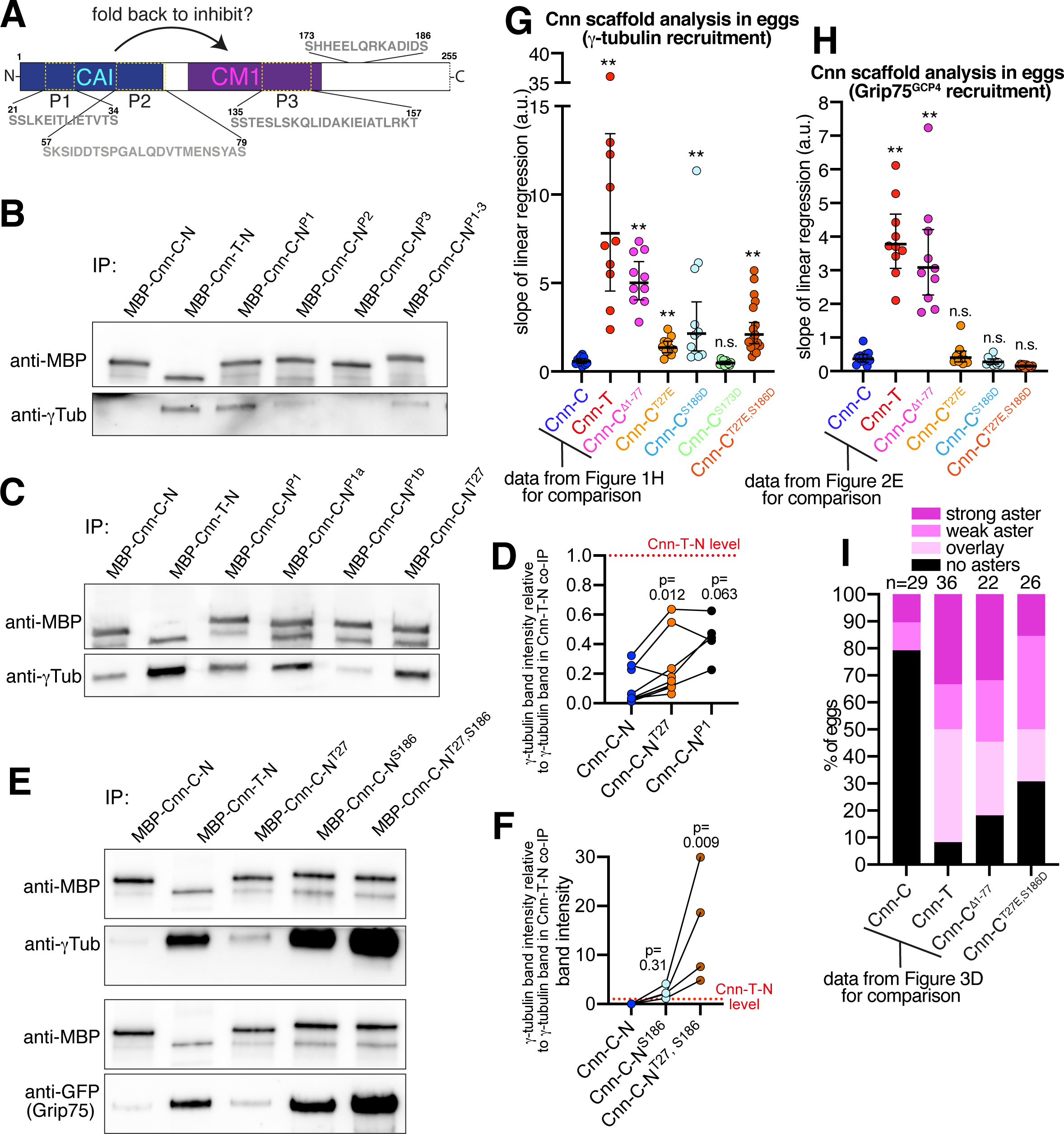
Phospho-mimetic mutations within the CAI domain and downstream of the CM1 domain promote binding to γ-tubulin complexes. **(A)** A cartoon showing the N-terminal region (aa1-255) of Cnn used in co-IP experiments. Regions of potential phosphorylation sites are indicated, with their amino acid sequence displayed. **(B-F)** Western blots of a co-IP experiments (B,C,E) and quantification of γ-tubulin bands (D,F) showing the efficiency to which different MBP-tagged N-terminal fragments of Cnn, as indicated, co-IP γ-tubulin from extracts of wild-type embryo (B-F), or γ-tubulin (top panels in E) and Grip75^GCP4^-sfGFP (bottom panels in E) from extracts of Grip75^GCP4^-sfGFP-expressing embryos. In (D) and (F), band intensities were normalised within each experiment to the γ-tubulin band in the respective MBP-Cnn-T-N IP. The connecting lines indicate data points obtained from within the same experiment. P values are from comparisons to the Cnn-C mean using either Wilcoxon matched-pairs signed rank tests (D, n=9 for comparison with Cnn-C-N^T27^; n=5 for comparison with Cnn-C-N^P1^) or a Dunn’s multiple comparisons test (F, n=4). (**G**,**H**) Graphs showing the S values from eggs expressing either γ-tubulin-mCherry (G) or Grip75^GCP4^-sfGFP (H) which contain the indicated scaffold types. Note that the data for Cnn-C, Cnn-T, and Cnn-C^Δ1-77^ scaffolds is the same as in Figures 1H and 2E to allow comparisons with the phospho-mimetic scaffolds. In (G) n= 2650 scaffolds and 11 eggs for Cnn-C^T27^ scaffolds, 1803 scaffolds and 11 eggs for Cnn-C^T186^ scaffolds, 2482 scaffolds and 10 eggs for Cnn-C^T173^ scaffolds, and 2835 scaffolds and 18 eggs for Cnn-C^T27,S186^ scaffolds. In (H) n= 1448 scaffolds and 10 eggs for Cnn-C^T27^ scaffolds, 1074 scaffolds and 10 eggs for Cnn-C^T186^ scaffolds, and 943 scaffolds and 10 eggs for Cnn-C^T27,S186^ scaffolds. The geometric mean and 95% CIs are indicated. ** indicates p<0.01, n.s. indicates p>0.05. P values were from comparisons to the Cnn-C mean using a one-way ANOVA of log_10_ transformed data. (**I**) Bar-graph showing results of a blind categorisation of eggs containing the different scaffold types based on the ability of the scaffolds within each egg to organise microtubule asters (numbers of eggs analysed indicated above). Note that the data for Cnn-C, Cnn-T, and Cnn-C^Δ1-77^ scaffolds is the same as in Figure 3D to allow comparisons with the phospho-mimetic scaffolds.

We therefore considered other putative phosphorylation sites. Phosphorylation slightly downstream of the CM1 domain promotes binding to γ-TuRCs in humans and *C. elegans* CM1 domain proteins (Ohta et al., 2021; Hanafusa et al., 2015). While the sequence surrounding the CM1 domain is not conserved across diverse species (Figure S3B), we identified two serine residues (S173 and S186) downstream of Cnn’s CM1 domain that were conserved in *Drosophila* species (Figure S3A). These sites also mapped to a similar predicted coiled-coil region to the sites in human CDK5RAP2 and *C. elegans’* SPD-5 (Figure S3B). While phospho-mimicking S^173^ had no effect, scaffolds with a phospho-mimic mutation at S^186^ (Cnn-C^S186D^ scaffolds) recruited ∼3.8-fold more γ-tubulin than Cnn-C scaffolds (Figure 4G). Moreover, N-terminal fragments containing this mutation (Cnn-C-N^S186^) co-IP’d γ-tubulin with a similar, if not higher, efficiency compared to the Cnn-T-N or Cnn-C^Δ1-77^ fragments (Figure 4E). In addition, although not apparent in the scaffold assay (Figure 4G), phospho-mimicking both T^27^ and S^186^ had a synergistic effect in the co-IP assay, where Cnn-C-N^T27,S186^ fragments co-IP’d significantly more γ-tubulin than any other type of fragment (Figure 4F). The same pattern was seen when co-immunoprecipitating the γ-TuRC-specific protein Grip75^GCP4^-sfGFP (Figure 4E). Unexpectedly, unlike in the co-IP assay, we did not see increased recruitment of Grip75^GCP4^-sfGFP to scaffolds containing any of the N-terminal phospho-mimetic mutations, including Cnn-C^T27,S186^ scaffolds (Figure 4H). This suggested that these scaffolds recruit γ-TuSCs rather than γ-TuRCs, potentially explaining why they do not recruit γ-tubulin to the levels seen at Cnn-T or Cnn-C^Δ1-77^ scaffolds (Figure 4G). Nevertheless, Cnn-C^T27,S168D^ scaffolds did organise microtubules more readily than Cnn-C scaffolds (Figure 4I – data compared to that in Figure 3D), suggesting that the γ-tubulin complexes bound by the phospho-mimetic forms of Cnn-C are at least semi-functional. Thus, while there are some differences between the scaffold assay and the co-IP assay, the data collectively suggest that phosphorylation at T^27^ and in particular at S^186D^ help to relieve CAI domain auto-inhibition and promote the binding of Cnn-C to γ-TuRCs.

### Ubiquitous expression of Cnn-C containing the high binding-affinity Cnn-T N-terminal region has a dominant negative effect and leads to fertility defects

We next wanted to test whether Cnn-C auto-inhibition is important for cell and developmental fidelity in *Drosophila*. We generated a transgenic fly line by random insertion of a ubiquitously-driven untagged Cnn-C construct in which its N-terminal region had been replaced with the N-terminal region of Cnn-T (pUbq-Cnn-C^T^) (Figure 5A). Based on our data so far, this form of Cnn should bind strongly to cytosolic γ-TuRCs but otherwise be regulated normally. We also generated a control line ubiquitously expressing untagged wild-type Cnn-C (pUbq-Cnn-C), whose binding to cytosolic γ-TuRCs should be restricted by the CAI domain.

**Figure 5.**
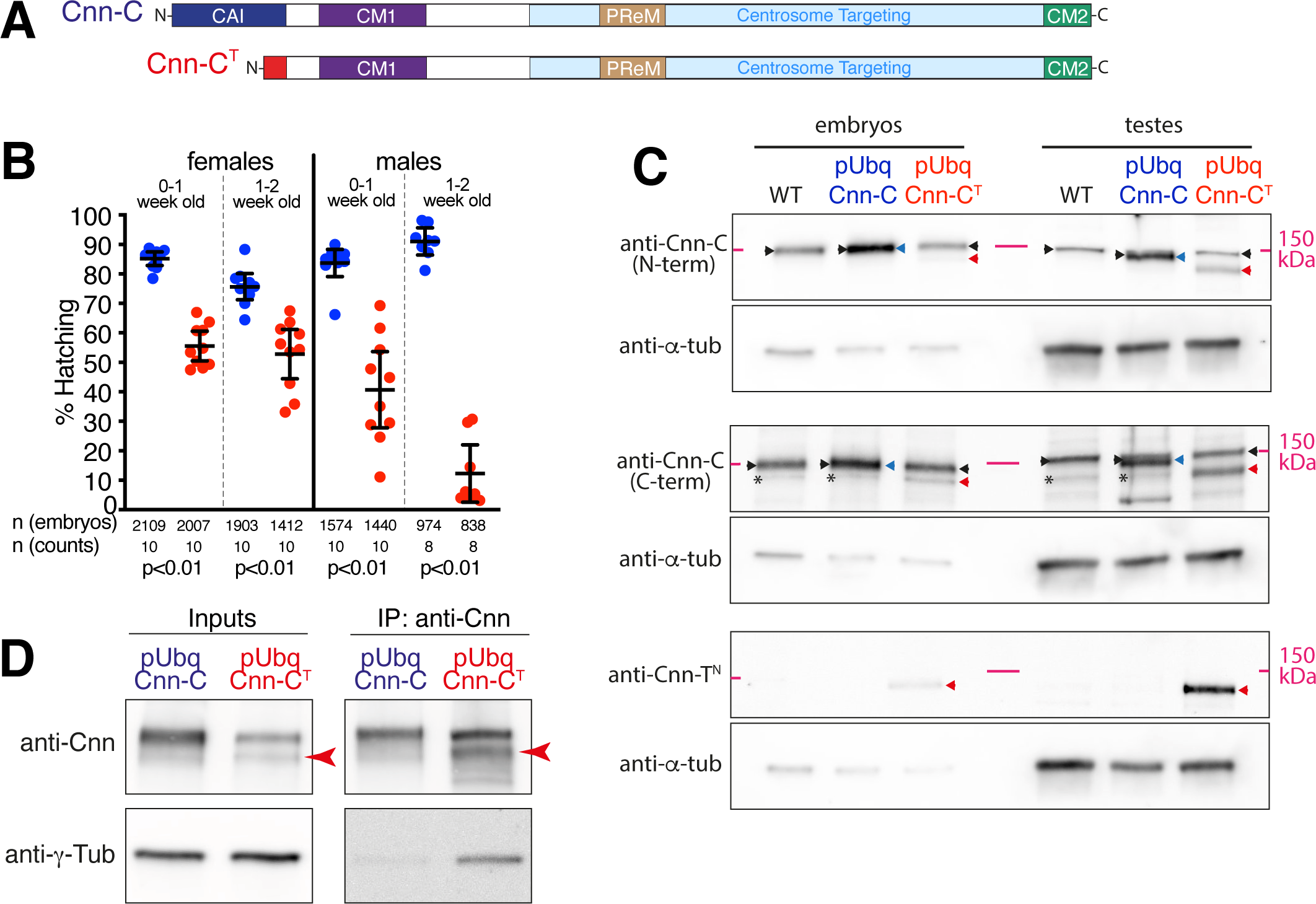
Expression of pUbq-Cnn-C^T^, which ectopically binds γ-TuRCs, reduces the ability of flies to generate progeny. **(A)** Diagram of normal Cnn-C and chimeric Cnn-C^T^ in which the CAI domain of Cnn-C (dark blue) is replaced by the shorter N-terminus of Cnn-T (red). **(B)** Graph showing the proportion of embryos that hatched from crosses of wild type flies to 0-1-or 1-2-week old pUbq-Cnn-C or pUbq-Cnn-C^T^ males or females, as indicated. Means and 95% confidence intervals are indicated. Total numbers of embryos counted and number of counts are indicated below. **(C)** Western blots of protein extracts from embryos and testes of wild-type (WT), pUbq-Cnn-C, and pUbq-Cnn-C^T^ flies, as indicated. Blots were probed with anti-γ-tubulin, anti-Cnn-C (N-term), anti-Cnn-C (C-term), and anti-Cnn-T^N^ antibodies as indicated. Note that endogenous Cnn-C (black arrowheads) runs at the same height as pUbq-Cnn-C (blue arrowheads) on these blots, explaining the increased brightness of these bands in the pUbq-Cnn-C extract lanes. Note also that the C-terminal Cnn-C antibody recognises an unspecific band (asterisks) of approximately the same size as pUbq-Cnn-C^T^ (red arrowheads) and thus the pUbq-Cnn-C^T^ band intensity would be lower in the absence of this unspecific band. **(D)** Western blot showing co-IP of γ-tubulin via anti-Cnn antibodies from embryo extracts expressing either pUbq-Cnn-C or pUbq-Cnn-C^T^, as indicated. Red arrowhead indicates Cnn-C^T^. Note that, given the low expression of pUbq-Cnn-C^T^ within embryos, gel loading of the IP lanes was adjusted to better balance the amount of Cnn protein per lane.

It was difficult to generate a viable pUbq-Cnn-C^T^ line and, once generated, was difficult to maintain and combine with other alleles. Thus, all following experiments were performed with the pUbq constructs expressed in the presence of endogenous Cnn. By crossing pUbq-Cnn-C and pUbq-Cnn-C^T^ females or males to wild-type flies and quantifying embryo hatching rates, we found that pUbq-Cnn-C^T^ flies were less able to generate progeny than pUbq-Cnn-C flies, with males being more affected than females (Figure 5B). Western blots of embryo or testes extracts using different Cnn-C antibodies and a Cnn-T-specific antibody showed that the level of pUbq-Cnn-C^T^ (red arrowheads) relative to endogenous Cnn-C (black arrowheads) was higher in testes extracts compared to embryo extracts (Figure 5C). In the embryo extracts, the pUbq-Cnn-C^T^ band was much weaker than the endogenous Cnn-C band, which is unusual for pUbq-driven Cnn constructs (P. Conduit unpublished observations), suggesting its expression was being suppressed. In contrast, the pUbq-Cnn-C^T^ band was of a similar intensity to, if not higher than, the endogenous Cnn-C band in the testes extracts. We therefore conclude that, relative to endogenous Cnn-C, pUbq-Cnn-C^T^ is weakly expressed within the maternal germline but is expressed to levels similar to endogenous Cnn within the testes. While other factors could be involved, such as cell-specific effects of Cnn to γ-TuRC binding, these differences in the expression levels of pUbq-Cnn-C^T^ between cells could explain the difference in the ability of male and female flies to generate progeny.

### Mis-regulation of binding to γ-tubulin complexes results in ectopic microtubule nucleation and defects during cell division

The failure of pUbq-Cnn-C^T^ flies to generate normal numbers of progeny suggested that ectopic binding of Cnn to γ-TuRCs leads to cellular defects during germline or early development. Co-IPs from embryo extracts confirmed that pUbq-Cnn-C^T^ binds γ-TuRCs more efficiently than pUbq-Cnn-C (Figure 5D). Moreover, binding of Cnn to γ-TuRCs appears to promote microtubule nucleation within the cytosol, as unfertilised eggs injected with mRNA encoding the GFP-tagged N-terminal region of Cnn-T, which efficiently binds γ-TuRCs, frequently displayed dynamic microtubules throughout their cytosol (9/12 eggs) (Video 5). This was not observed in all 27 control-injected eggs. This effect is similar to that observed when expressing CM1-domain fragments within human and fission yeast cells (Choi et al., 2010; Cota et al., 2017; Hanafusa et al., 2015; Lynch et al., 2014) and suggests that CM1 domain binding to γ-TuRC also promotes microtubule nucleation in *Drosophila*.

To examine whether ectopic cytosolic microtubules formed within cells expressing pUbq-Cnn-C^T^ and whether there were associated cell defects, we immunostained various fly tissues. There were no obvious defects within oocytes from pUbq-Cnn-C^T^ flies, where the positions of the nucleus and Gurken protein, both dependent on proper microtubule organisation, were normal in oocytes from both pUbq-Cnn-C and pUbq-Cnn-C^T^ females (Figure S4A-C). We did, however, frequently observe defects in fixed and stained syncytial embryos from pUbq-Cnn-C^T^ females (Figure 6D-F) as compared to embryos from pUbq-Cnn-C females (Figure 6A-C). These defects included: an apparent excess of cytosolic microtubules, unusually bright microtubule asters, and nuclear organisation defects during S-phase (Figure 6D); highly disorganised spindles during M-phase (Figure 6E); and an apparent excess of cytosolic microtubules during telophase (Figure 6F). In a blind analysis of embryos, severe and moderate defects were observed in a higher proportion of embryos from pUbq-Cnn-C^T^ females (19.4% severe and 30.6% moderate) than from wild-type (4.1% and 18.4%) or pUbq-Cnn-C (3.4% and 25%) females (Figure 6G). While half of the embryos from pUbq-Cnn-C^T^ females were normal, this could reflect the relatively low expression of pUbq-Cnn-C^T^ in the female germline (Figure 5C). Broadly, the categorisation of embryo defects in Figure 6G reflects the observed hatching rates in Figure 5B, assuming that embryos with moderate and severe defects often fail in development.

**Figure 6.**
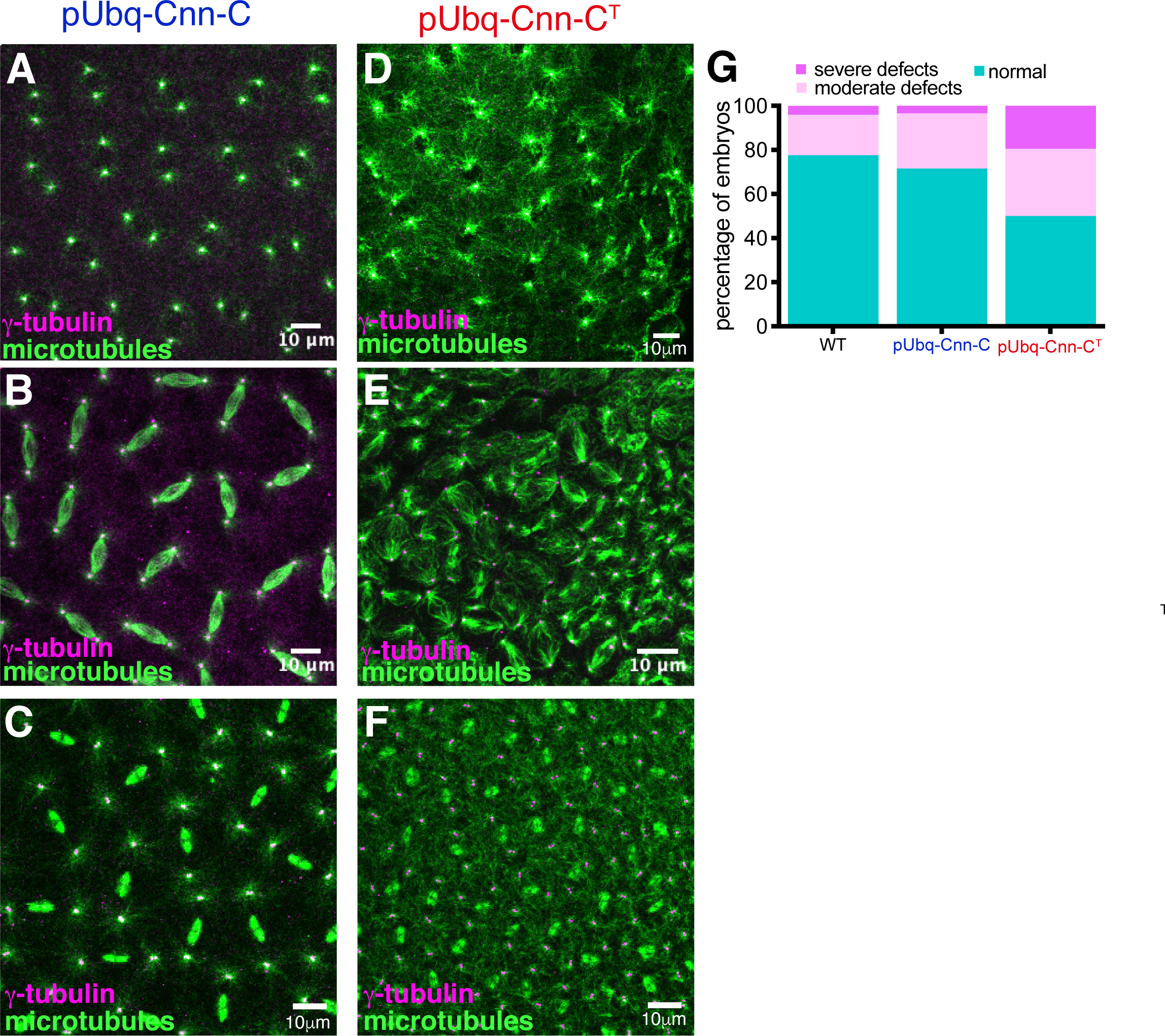
Expression of pUbq-Cnn-C^T^ increases the frequency of nuclear and spindle defects observed within syncytial embryos. **(A-F)** Fluorescence images of syncytial embryos expressing either pUbq-Cnn-C (A-C) or pUbq-Cnn-C^T^ (D-F) in either S-phase/prophase (A,D) Metaphase (B,E), or telophase (C,F). Note the apparent high density of cytosolic microtubules that can be (but are not always) observed in pUbq-Cnn-C^T^ embryos, along with major organisation defects. **(G)** Graph showing results from a blind categorisation of wild-type (n=49), pUbq-Cnn-C (n=88), or pUbq-Cnn-C^T^ (n=36) embryos based on the presence or absence of moderate or severe nuclear or spindle defects.

Consistent with a very strong reduction in the ability of pUbq-Cnn-C^T^ males to generate progeny, defects were frequently observed within their testes, where production of sperm involves a series of mitotic and meiotic cell divisions. When meiosis progresses normally, the 64 round spermatids cells within the resulting cyst all contain a similarly sized phase-light nucleus and phase-dark nebenkern (which is an accumulation of mitochondria that were segregated during meiosis). This was true in round spermatids from pUbq-Cnn-C testes (Figure 7A,C), but not in round spermatids from pUbq-Cnn-C^T^ testes (Figure 7B,C), suggesting that pUbq-Cnn-C^T^ expression results in problems in chromosome segregation and cytokinesis. Indeed, a high density of cytosolic microtubules and clear meiotic defects were observed in spermatocytes within fixed and stained pUbq-Cnn-C^T^, but not pUbq-Cnn-C, testes. Defects were observed at various developmental stages and included cells with incorrect numbers of nuclei and centrosomes as well as cells containing multiple spindles (Figure 7D,E; Figure S5A,B). Thus, ectopic binding of pUbq-Cnn-C^T^ to γ-TuRCs within spermatocytes appears to lead to ectopic microtubule nucleation within the cytosol and major defects during meiosis.

**Figure 7.**
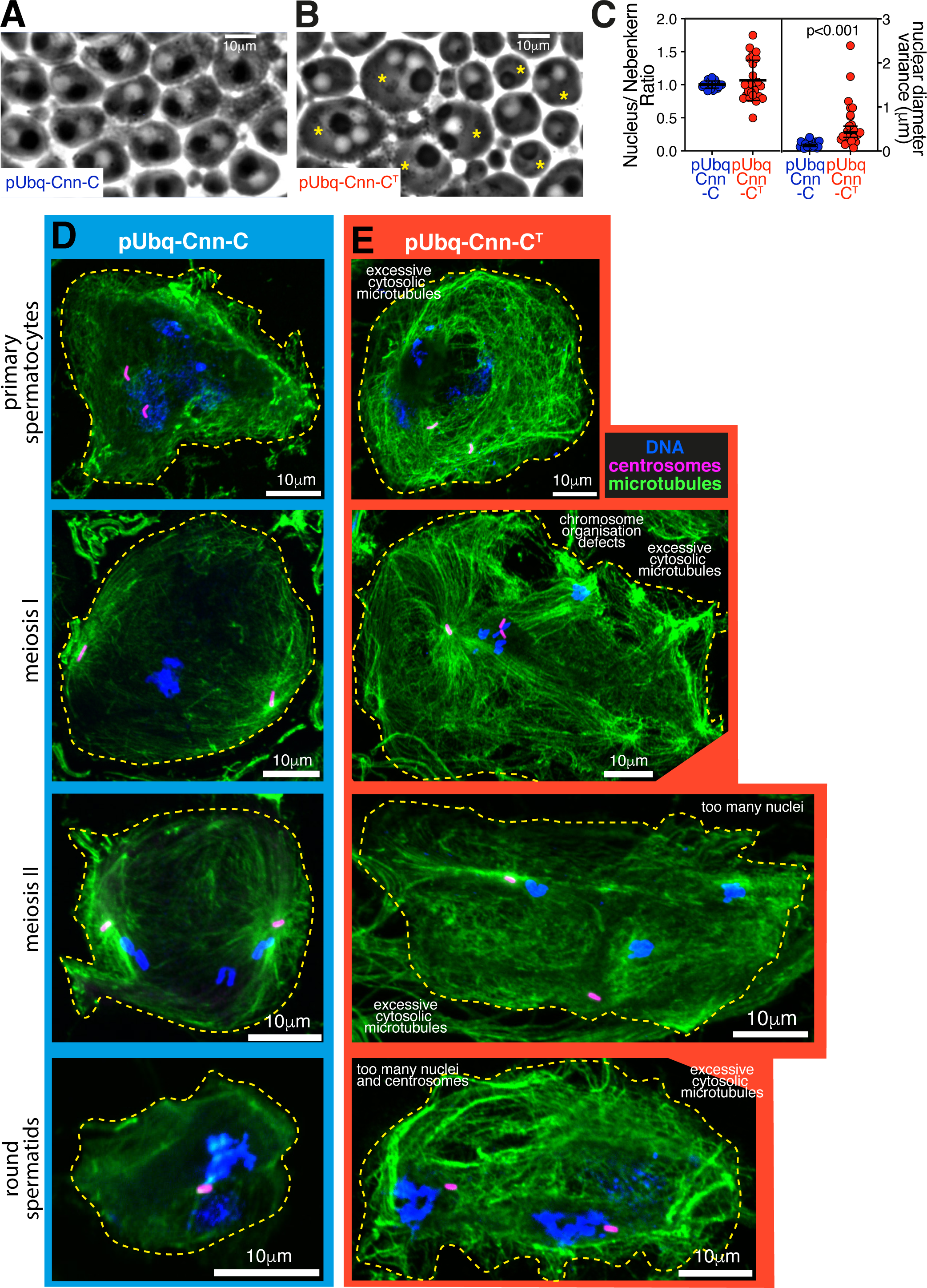
Expression of pUbq-Cnn-C^T^ results in major defects during male meiosis. **(A,B)** Phase contrast images showing round spermatids from testes of flies expressing pUbq-Cnn-C (A) or pUbq-Cnn-C^T^ (B). Alterations in nucleus: nebenkern ratio (normally 1:1, asterisks in right panel) and size (normally approximately equal) indicate defects in cytokinesis and karyokinesis. **(C)** Graph showing quantification of the nucleus:nebenkern ratio (left panel - means and standard deviations indicated) and variance in nuclear diameter (right panel – geometric means and 95% CIs indicated, p value from an unpaired t-test of log_10_-tranformed data) in pUbq-Cnn-C (n=22 cysts) and pUbq-Cnn-C^T^ (n=27 cysts) testes. **(D**,**E)** Fluorescence images showing spermatocytes or round spermatids at different developmental stages, as indicated, from testes of flies expressing pUbq-Cnn-C (D) or pUbq-Cnn-C^T^ (E) stained for microtubules (green, *α*-tubulin), centrosomes (pink, Asterless), and DNA (blue). Defects within cells expressing pUbq-Cnn-C^T^ include an apparent high density of cytosolic microtubules, abnormal spindles, and too many nuclei.

### Human CDK5RAP2 binding to γ-TuRCs is also regulated by auto-inhibition, but the precise mechanism differs from *Drosophila*

To examine whether auto-inhibition is a conserved feature of CM1 domain proteins, we tested the ability of various N-terminal fragments of human CDK5RAP2 (Figure 8A) to co-IP γ-tubulin from HEK cell extracts. The reported CM1 domain spans aa58-126 of CDK5RAP2 (Sawin et al., 2004; Zhang and Megraw, 2007) (Figure 8A) and a fragment spanning aa1-210 was less efficient at co-IP’ing γ-tubulin than a fragment spanning aa51-100 (also known as γ-TuNA (Choi et al., 2010)) (Figure 8A,B). This indicated that sequences either upstream or downstream of γ-TuNA are inhibitory for binding to γ-TuRCs. A fragment that included the sequence upstream of γ-TuNA (aa1-100) co-IP’d γ-tubulin more efficiently than γ-TuNA (Figure 8B), suggesting that, unlike in *Drosophila* Cnn, the sequence upstream of the CM1 domain is not inhibitory but is instead required for efficient binding. In contrast, a fragment that included the sequence downstream of γ-TuNA (aa51-210) was less efficient than γ-TuNA at co-IP’ing γ-tubulin (Figure 8B). This suggests that the sequence downstream of CDK5RAP2’s CM1 domain inhibits binding to γ-TuRCs. Thus, while auto-inhibition appears to regulate the binding of CDK5RAP2 to γ-TuRCs as in flies, the precise mechanism appears to vary between species.

**Figure 8.**
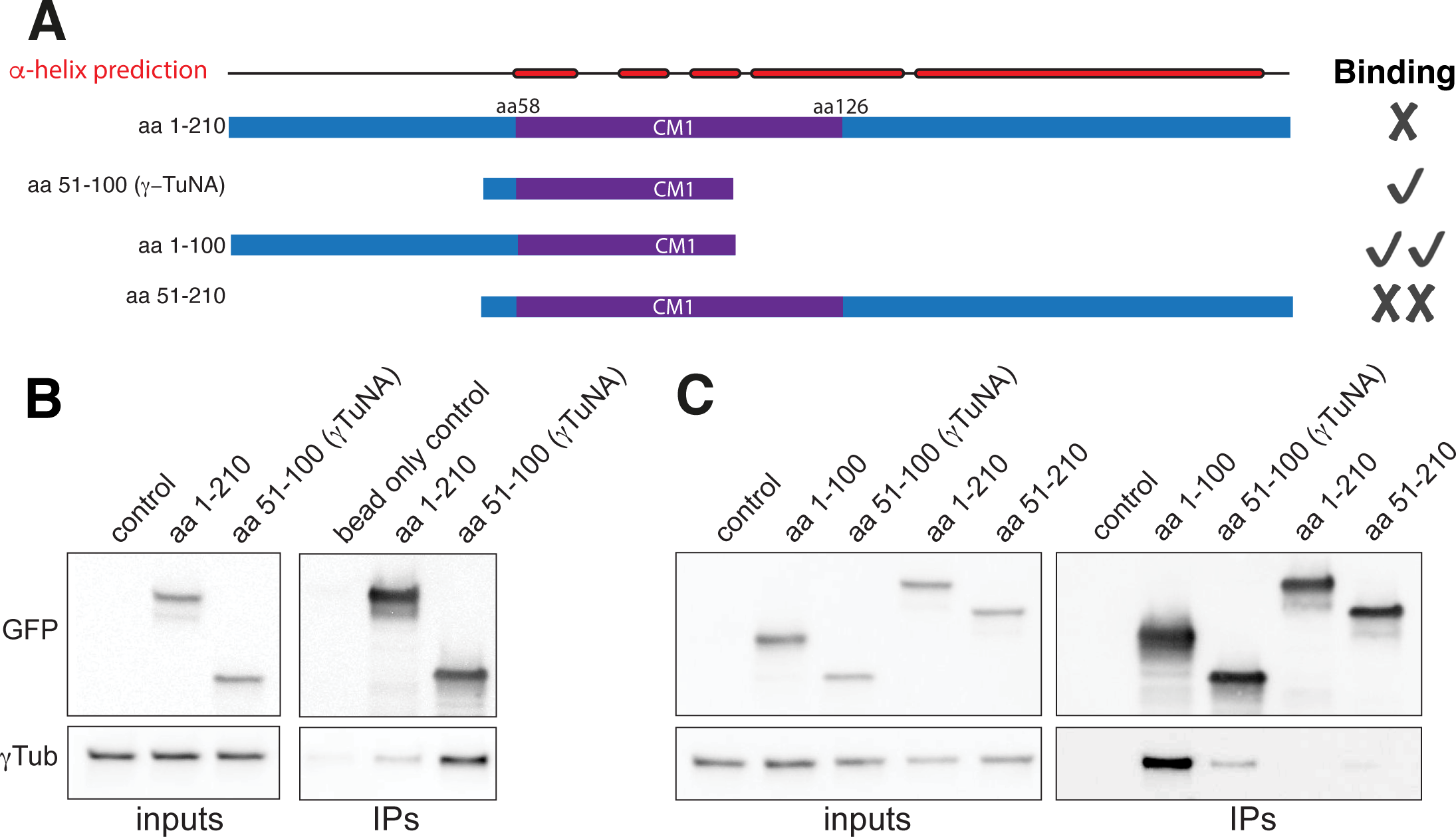
The region downstream of the CM1 domain in human CDK5RAP2 is inhibitory for binding to γ-TuRCs. **(A)** Cartoon depicting the various CDK5RAP2 N-terminal fragments used in IP experiments and indicating their relative γ-TuRC binding affinity. **(B,C)** Western blots of co-IP experiments from HEK cell extracts probed for the various GFP-tagged CDK5RAP2 fragments (top) and γ-tubulin (bottom).

## Discussion

We have shown that the extreme N-terminal region of Cnn-C, which we name the CAI domain, inhibits Cnn-C from binding to γ-TuRCs. This auto-inhibition is important because expressing a form of Cnn that readily binds γ-TuRCs within the cytosol leads to spindle and cell division defects, possibly via the ectopic activation of γ-TuRCs. Phospho-mimicking experiments suggest that phosphorylation at sites close to the CM1 domain relieves auto-inhibition of Cnn-C and promotes binding to γ-TuRCs. This is consistent with Cnn-C being phosphorylated specifically at centrosomes during mitosis (Conduit et al., 2014a) where binding and activation of γ-TuRCs takes place. In addition, human CDK5RAP2 is inhibited from binding cytosolic γ-TuRCs by the region downstream of the CM1 domain. Thus, while the precise mechanism may vary, it appears that auto-inhibition is a conserved feature of CM1 domain proteins.

There is considerable evidence, including the work presented here, showing that binding of CM1-domain proteins to γ-tubulin complexes stimulates microtubule nucleation (Choi et al., 2010; Muroyama et al., 2016; Hanafusa et al., 2015; Cota et al., 2017; Lynch et al., 2014), but the reason remains unclear. One possibility is that binding leads to conformational changes in γ-TuRCs, but human and *Xenopus* γ-TuRCs bound by CM1 domain fragments remain in an open, seemingly inactive, conformation (Wieczorek et al., 2019; Liu et al., 2019). Whether this is due to a low stoichiometry of binding remains unclear, but binding of the CM1 domain to *S. cerevisiae* γ-TuSCs/γ-TuRCs does result in structural changes that possibly promote nucleation activity (Brilot et al., 2019). It is also possible that CM1 domain binding has a context-specific effect. Adding CM1-domain fragments to purified γ-TuRCs within *Xenopus* egg extracts had a greater effect on nucleation efficiency when the extract was supplemented with activated Ran (Liu et al., 2019), and we find that expression of pUbq-Cnn-C^T^ leads to defects within specific cell types (although these differences could simply be due to the differences in expression levels). Clearly, we need a better understanding of how CM1 domain binding promotes microtubule nucleation.

Phosphorylation appears to be an important mechanism for promoting binding between CM1 domain proteins and γ-TuRCs. This is true for human CDK5RAP2, *C. elegans* SPD-5, and *S. cerevisiae* SPC110, where the phosphorylation sites that promote binding have been identified either upstream or downstream of the CM1 domain (Hanafusa et al., 2015; Lin et al., 2014b; Ohta et al., 2021). We show that phospho-mimicking sites both upstream and downstream of the CM1 domain also promotes binding of *Drosophila* Cnn-C to γ-TuRCs. We predict that phosphorylation helps to relieve the auto-inhibition imposed by the CAI domain as well as directly increasing binding affinity between Cnn-C and γ-TuRCs. We find that phospho-mimicking S^186^ alone allows robust binding to γ-TuRCs, suggesting that phosphorylating this single site is sufficient to relieve auto-inhibition fully, at least *in vitro*. Phospho-mimicking T^27^ has a more subtle effect but has a strong effect when S^186^ is also phospho-mimicked. This suggests that phospho-mimicking T^27^ increases the binding affinity between Cnn-C and γ-TuRCs, rather than relieving auto-inhibition, and thus has a minimal effect when Cnn-C is auto-inhibited (when S^186^ is not phospho-mimicked) but has a strong effect when Cnn-C inhibition is relieved (by S^186^ phospho-mimicking). This would suggest that the CAI domain, which contains T^27^, is also involved in binding to γ-TuRCs once inhibition is relieved. A role for the region upstream of the CM1 domain in binding to γ-TuRCs may be conserved, as our data shows this region is promotes binding of human CDK5RAP2 to γ-TuRCs.

In future, it will be important to understand how the CAI domain inhibits the CM1 domain. We previously postulated that the extreme N-terminal region of Cnn-C (i.e. the CAI domain) might fold back and sterically inhibit the CM1 domain (Tovey et al., 2018). Our data is consistent with this possibility and, in our view, this is the most likely explanation. A similar mechanism has also been proposed in *C. elegans* (Ohta et al., 2021). Nevertheless, there are alternative possibilities, including that the CAI domain could recruit another protein that interferes with CM1 domain binding. In any case, it will be interesting to compare how auto-inhibition is achieved in different homologues, especially given that the region downstream, not upstream, of the CM1 domain appears to mediate inhibition in human CDK5RAP2.

Importantly, our data also highlights differences in how binding between CM1 domain proteins and γ-TuRCs is regulated within different cell types and at different MTOCs. We have shown that the testes specific Cnn-T isoform, which lacks the CAI domain, can bind efficiently to γ-tubulin complexes in the apparent absence of any upstream regulatory events. Cnn-T is expressed primarily within developing sperm cells and isoform-specific C-terminal exons mediate its recruitment to mitochondria, where it binds and recruits γ-tubulin complexes (Chen et al., 2017). The mitochondrial surface is very different from mature centrosomes, which concentrate a selection of kinases. It therefore seems appropriate that Cnn-T isoforms do not appear to rely on phosphorylation for binding. Presumably, binding and potential activation of γ-TuRCs within the shrinking cytosol of developing sperm cells is not detrimental to sperm development (and may even be important for amplifying cytoplasmic microtubules), unlike in dividing cells where our data shows that spindle formation and cytokinesis are clearly perturbed.

In summary, the data presented here provide important insights into how and why binding of CM1 domain proteins to γ-TuRCs is regulated. Future studies will help elucidate the precise mechanism underlying auto-inhibition of the CM1 domain and how this may vary between species. It will also be important to determine whether CM1 domain binding directly activates γ-TuRCs and, if not, how CM1 domain binding promotes microtubule nucleation.

## Materials and Methods

### DNA cloning

5-alpha competent *E. coli* cells (high efficiency, NEB) were used for bacterial transformations. DNA fragments were purified using QIAquick Gel Extraction kits (Qiagen); plasmid purification was performed using QIAprep Spin Miniprep kits (Qiagen). Phusion high-fidelity PCR master mix with HF buffer (ThermoFisher Scientific) was used for PCRs.

### Transgenic *Drosophila* lines

All endogenously-tagged lines were made using CRISPR combined with homologous recombination, by combining the presence of a homology-repair vector containing the desired insert with the appropriate guide RNAs and Cas9. The γ-tubulin37C-mCherry and Grip128-sfGFP alleles were generated by inDroso. For γ-tubulin37C-mCherry, eggs from nos-Cas9 expressing females were co-injected with a plasmid encoding the expression of dual guides targeting each side of the 3’UTR, TACACATATCAAGATACATG and CCCAGATCGATTATCCCCAG, and a plasmid containing a SSSS-mCherry-3’UTR-LoxP-3xP3-dsRED-Lox P cassette flanked by homology arms (the multi-serine insert acts as a flexible linker). After screening for dsRED, the selection marker was excised by Cre recombination. For Grip128-sfGFP, eggs from nos-Cas9 expressing females were co-injected with a plasmid encoding the expression of a single guide containing the target sequence ATGGGGCACACTGGAGTTGA and with a pBluescript plasmid containing sfGFP and linker sequence (4X GlyGlySer) flanked on either side by 1.5kb of DNA homologous to the genomic locus surrounding the 3’ end of the appropriate coding region. The homology vector was made within the lab (and sent to InDroso) by HiFi assembly (NEB) of PCR fragments generated from genomic DNA prepared from nos-Cas9 flies (using MicroLYSIS, Microzone) and a vector containing the sfGFP tag (DGRC, 1314). Screening for the insert was performed with the following primers: AGGAAGATGCGAACACACGT and GTACAGCTCATCCATGCCCA.

The Grip75-sfGFP and Grip163-sfGFP lines were made within the lab following a similar approach to that used previously (Tovey et al., 2018; Mukherjee et al., 2020). Flies expressing a single guide RNA containing the target sequence CAAAAACATCGTATTCATG or ACCACTATTACAAGGTATCT for Grip75-sfGFP or Grip163-sfGFP, respectively, were crossed to nos-Cas9 expressing females and the resulting embryos were injected with homology vectors by the Department of Genetics Fly Facility, Cambridge, UK. The homology vectors comprised a pBluescript plasmid containing sfGFP and linker sequence (4X GlyGlySer) flanked on either side by 1.5kb of DNA homologous to the genomic locus surrounding the 3’ end of the appropriate coding region. The homology vectors were made as for Grip128-sfGFP. F1 and F2 males were screened by PCR using the following primers: for Grip75-sfGFP: GAGAAGTTTGCGCATATGACCC and AGCAGCACCATGTGATCGCGC; for Grip163-sfGFP: AGTCGCAGTCCTTTATTGTGG and AGCAGCACCATGTGATCGCGC.

pUbq-Cnn-C and pUbq-Cnn-C^T^ were made from a pDONR-Cnn-C vector (gift from Jordan Raff). To generate a Cnn-T-specific N-terminal region of Cnn, an appropriate DNA fragment (made by GENEWIZ, based on the FlyBase sequence of Cnn-T) was synthesised and amplified by PCR and used to replace the N-terminal region of Cnn in a pDONR-Cnn-C vector cut with XmaI. The pDONR-Cnn-C and newly made pDONR-Cnn-T vectors were then inserted into a pUbq transformation vector (gift from Jordan Raff) by Gateway cloning (ThermoFisher Scientific). All DNA vectors were injected into embryos by the Department of Genetics Fly Facility, Cambridge, UK.

The Jupiter-mCherry line used to monitor microtubule nucleation was a gift from Jordan Raff’s lab. The original line was a GFP trap line from Daniel St. Johnston’s lab and the GFP was replaced with mCherry.

### Recombinant protein cloning, expression and purification

Fragments of Cnn-C-N and Cnn-T-N used in co-IP experiments were amplified from the pDONR-Cnn-C and pDONR-Cnn-T vectors described above by PCR and inserted into a pDEST-HisMBP (Addgene, #11085) vector by Gateway cloning (Thermo Fisher Scientific). Proteins were expressed in *E. coli* (BL21-DE3) and purified using affinity chromatography. MBP-tagged fragments were purified by gravity flow through amylose resin (New England Biolabs) and step elution in maltose. The concentration of each fraction was determined on a Nanodrop and peak fractions were diluted 1:1 with glycerol and stored at -20°C. Truncated fragments of Cnn-C were made by modification of the pDONR-Cnn-C-N entry clone. The N-terminal region was removed by a Quikchange reaction (Agilent), and the resulting shortened fragment was inserted into the pDEST-HisMBP destination vector via a Gateway reaction.

Phospho-mimetic fragments were created by modifying the pDONR-Cnn-C-N entry clone. The pDONR-Cnn-C-N backbone was linearised by PCR or by digestion, omitting the phospho-patch to be replaced. Phosphomimetic patches in which all S/T residues were swapped for D/E residues, respectively, were synthesised either by PCR using two overlapping primers or by GENEWIZ. They were inserted into the linear backbone by HiFi Assembly (NEB). The entry clones were checked by restriction enzyme digest and sequencing before being inserted into the pDEST-HisMBP destination vector via a Gateway reaction.

pRNA vectors were made by modification of the pDONR-Cnn-C-PReM^P^ vector containing phospho-mimetic mutations in the PReM domain (Conduit et al., 2014a). N-terminal variants were introduced by restriction digests (SspI-HF and AatII) of pDONR-Cnn-C, pDONR-Cnn-T, and pDONR-Cnn-C-PReM^P^ entry clones. Fragments were combined as necessary by NEB HiFi assembly to create new pDONR vectors, which were inserted into a pRNA-GFP or pRNA-mKate destination vector (Conduit et al., 2014a) via a Gateway reaction. The Cnn-T-N fragment was inserted directly into pRNA-GFP destination vectors via Gateway cloning.

Fragments of CDK5RAP2 were synthesised by Genewiz, amplified by PCR, and cloned into a pCMV-GFP vector (gift from Jens Lüders) by restriction digest and HiFi assembly (NEB).

### Primers

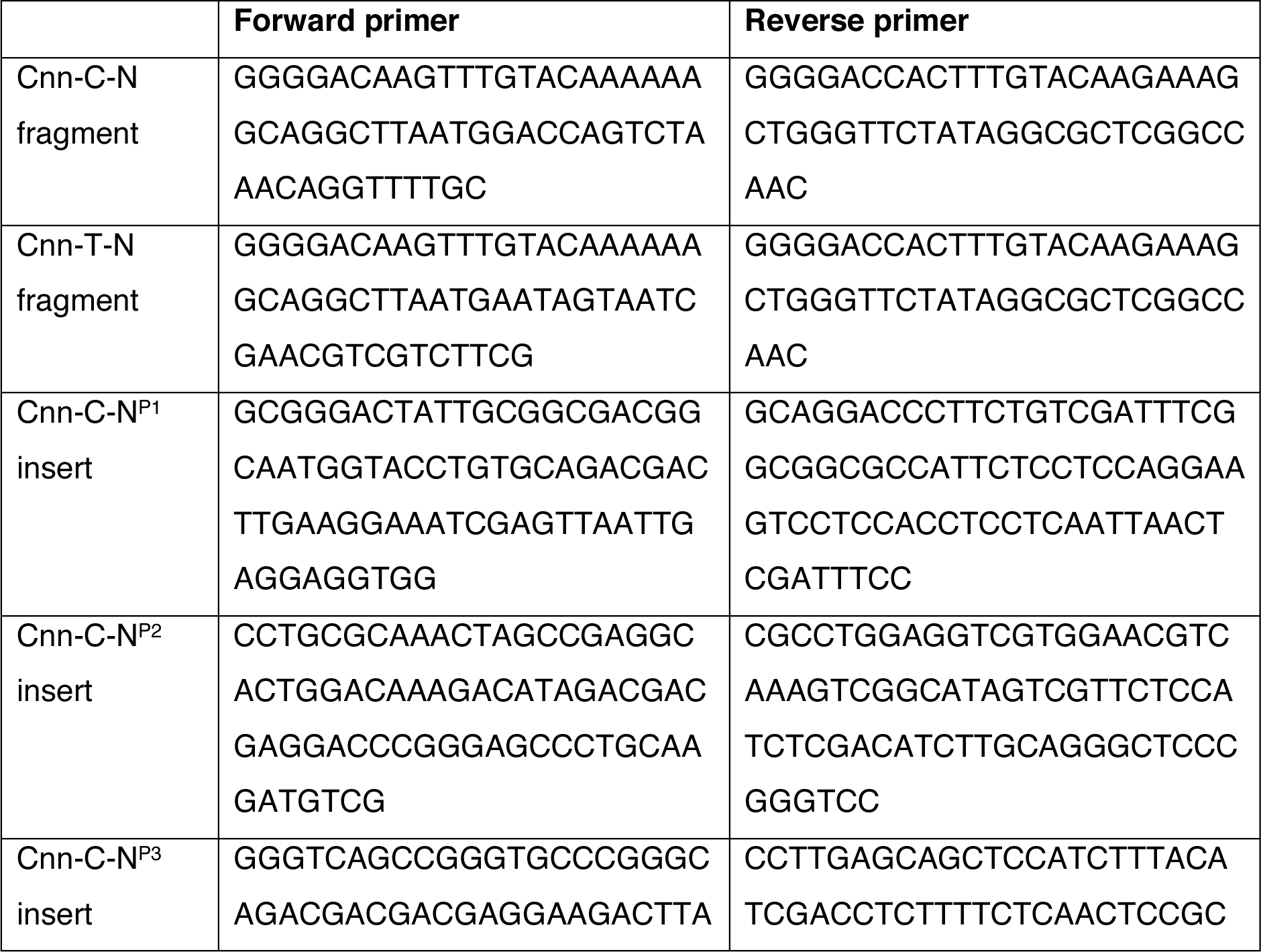
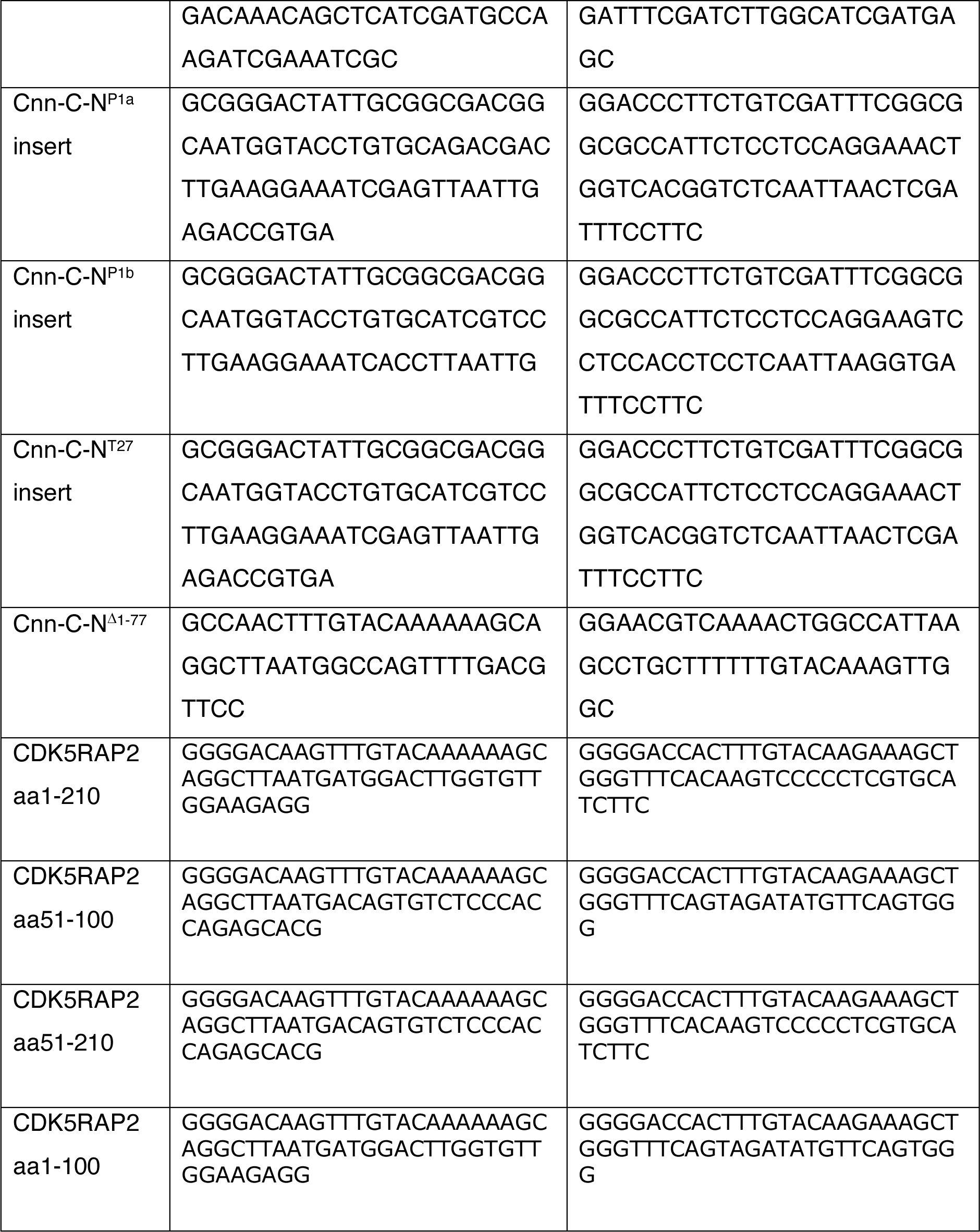

### Immunoprecipitation

1g/ml of embryos were homogenised with a hand-pestle in homogenisation buffer containing 50 mM HEPES, pH7.6, 1mM MgCl_2_, 1 mM EGTA, 50 mM KCl supplemented with PMSF 1:100, Protease Inhibitor Cocktail (1:100, Sigma Aldrich) and DTT (1M, 1:1000). Extracts were clarified by centrifugation twice for 15 minutes at 16,000 rcf at 4°C.

For the MBP-Cnn fragment IPs, 30 μl magnetic ProteinA dynabeads (Life Technologies) coupled to anti-MBP antibodies (gift from Jordan Raff) were incubated with an excess of purified MBP-Cnn fragments and rotated for 1 hour at 4°C. Unbound fragments were washed off in PBST, and the saturated beads were resuspended in 100 μl embryo extract and rotated at 4°C overnight. Beads were washed 5 times for 1 min each in PBST, boiled in 50 μl 2x sample buffer, and separated from the sample using a magnet. Samples were analysed by western blotting as described.

For the Grip-GFP IPs, 20 μl high-capacity ProteinA beads (Abcam) coupled to anti-MBP antibodies (gift from Jordan Raff) were incubated with an excess of purified MBP-Cnn fragments and rotated at 4°C for 1 hour. Unbound fragments were washed off in PBST and the saturated beads were resuspended in 65 μl embryo extract and rotated at 4°C overnight. Beads were washed 5 times for 1 min each in PBST, boiled in 2x sample buffer, and separated from the sample by centrifugation. Samples were analysed by western blotting as described.

For the IPs from pUbq-Cnn-C and pUbq-Cnn-C^T^ embryo extract, 50 μl magnetic ProteinA dynabeads (Life Technologies) coupled to anti-Cnn (C-terminal) antibodies (gift from Jordan Raff) were rotated in 100 μl embryo extract at 4°C overnight. Beads were washed 5 times for 1 min each in PBST, boiled in 2x sample buffer, and separated from the sample using a magnet. Samples were analysed by western blotting as described. We had tried these IPs using beads coated with the anti-Cnn-T^N^ antibody but found that they did not pull down any protein (data not shown), presumably as this antibody was raised against a peptide antigen and recognises only denatured pUbq-Cnn-C^T^ on western blots.

### Electrophoresis and western blotting

Samples were run on 4-20% TGX Precast Gels (BioRad) (except Figure 5C and D, in which samples were run on 7.5% TGX Precast gels (BioRad)), alongside 5μl Precision Plus WesternC Standard markers (BioRad). For western blotting, semi-dry blotting was carried out using TransBlot Turbo 0.2μm nitrocellulose membrane transfer packs (BioRad), and a TransBlot Turbo transfer system running at 1.3A, up to 25V, for 7 minutes (BioRad mixed molecular weight pre-set programme). Membranes were stained with Ponceau and washed, first with distilled water then with milk solution (PSBT + 4% milk powder), and then blocked in milk solution for 1 hour at room temperature. Sections of blots were incubated with primary antibodies as indicated in figures (antibodies found in table). Blots were incubated with horseradish peroxidase (HRP)-conjugated anti-mouse, anti-rabbit, or anti-sheep secondary antibodies (1:2000 in PSBT + 4% milk powder, ImmunoReagents) as appropriate for 45 mins at room temperature, washed in PSBT 3 times for 15 mins each, and then incubated with ECL substrate (BioRad ECL Clarity or ThermoFisher SuperSignal West Femto Max) for 5 minutes. Membranes were imaged using a Kodak Image Station 4000R or a BioRad ChemiDoc.

### Mass spectrometry

Samples were run into TGX Precast Gels (BioRad) and the gels were rinsed in dH_2_O. Bands were excised using a clean razor blade and cut into 1mm^2^ pieces on a fresh glass slide and placed into a microtube. Co-IP samples were processed by the Mass Spectrometry facility at the Department of Biochemistry, University of Cambridge with LC-MS/MS analysis using a Dionex Ultimate 3000 RSLC nanoUPLC (Thermo Fisher Scientific Inc, Waltham, MA, USA) system and a Q Exactive Orbitrap mass spectrometer (Thermo Fisher Scientific Inc, Waltham, MA, USA).

Post-run, all MS/MS data were converted to mgf files and the files were then submitted to the Mascot search algorithm (Matrix Science, London UK, version 2.6.0) and searched against the Uniprot Drosophila_melanogaster_20180813 database (23297 sequences; 16110808 residues) and common contaminant sequences containing non-specific proteins such as keratins and trypsin (123 sequences; 40594 residues). Variable modifications of oxidation (M), deamidation (NQ) and phosphorylation (S,T and Y) were applied as well a fixed modification of carbamidomethyl (C). The peptide and fragment mass tolerances were set to 20ppm and 0.1 Da, respectively. A significance threshold value of p<0.05 and a peptide cut-off score of 20 were also applied.

### Antibodies

Primary antibodies used in the study are indicated in the table below. For western blotting, primary and secondary antibodies were diluted in PBST + 4% milk; primary antibodies were diluted at concentrations indicated in the table; secondary antibodies were diluted at 1:2000. For immunostaining, primary and secondary antibodies were diluted in PBS + 0.1% Triton (PBT) + 5% BSA; primary antibodies were diluted at concentrations indicated in the table; secondary antibodies (AlexaFluor 488, 561, or 633 conjugated secondary antibodies (ThermoFisher)) were diluted at 1:1000 for testes and 1:1500 for embryos. DNA was stained with Hoechst (Life Technologies, 33342) or DAPI.

| Antibody | WB Concentration | IF Concentration | Source |
| --- | --- | --- | --- |
| α-Tubulin mouse monoclonal | - | 1:1000 | Sigma Aldrich, DM1a |
| Asl (N-terminal) guinea pig polyclonal | 1:1000 | 1:1000 | Gift from Jordan Raff |
| Cnn (N-terminal) rabbit monoclonal | 1:1000 | 1:1000 | Gift from Jordan Raff |
| Cnn (C-terminal) sheep polyclonal | 1:1000 | - | Gift from Jordan Raff |
| Cnn-T<sup>N</sup> Rabbit polyclonal | 1:500 | - | This study |
| γ-Tubulin mouse monoclonal | 1:500 | 1:500 | Sigma Aldrich, GTU-88 |
| γ-Tubulin rabbit polyclonal | - | 1:500 | Sigma Aldrich, T5192 |
| GFP mouse monoclonal | 1:250 or 1:500 | 1:250 or 1:500 | Roche, 11814460001 |
| Grip71 rabbit polyclonal | 1:100 | 1:100 | CRB (crb2005268) |
| MBP rabbit polyclonal | 1:3000 | - | Gift from Jordan Raff |
| Phospho-histone H3 rabbit polyclonal | - | 1:500 | Abcam, AB5176 |
| Staufen Mouse monoclonal | - | 1:100 | Santa Cruz dN-16 |
| Gurken Mouse monoclonal | - | 1:200 | DSHB 1D12 |
| Lamin Dm0 | - | 1:30 | DSHB 84.12 |

### Immunostaining

Testes were dissected in PBS, fixed in 4% paraformaldehyde for 30 minutes, washed 3x 5 minutes in PBS and incubated in 45% and then 60% acetic acid before being squashed onto slides and flash-frozen in liquid nitrogen. Coverslips were removed and samples were post-fixed in methanol at -20°C, washed 3x 15 minutes in PBS + 0.1% Triton (PBST), then incubated overnight in a humid chamber at 4°C with primary antibodies diluted in PBST + 5% BSA + 0.02% azide. Slides were washed 3x 5 minutes in PBST and then incubated for 2 hours at room temperature with Alexa-Fluor secondary antibodies (ThermoFisher) (all 1:1000 in PBST + 5% BSA + 0.02% azide). Slides were washed 3x 15 minutes in PBST, 10 minutes in PBST with Hoechst, and then 5 minutes in PBST. 10 μl of mounting medium (85% glycerol in water + 2.5% N-propyl-galate) was placed on top of the tissue and a coverslip was gently lowered and sealed with nail varnish.

Embryos were collected within 2-3 hours of laying and were dechorionated in 60% bleach for 2 minutes. Vitelline membranes were punctured with a combination heptane and methanol + 3% EGTA (0.5M) before three washes in neat methanol. Embryos were fixed in methanol at 4°C for at least 24 hours before rehydrating. Embryos were rehydrated by washing 3x 20 mins in PBST, then blocked in PBST + 5% BSA for 1 hour, followed by overnight incubation in primary antibodies in PBST + 5% BSA at 4°C. Embryos were washed 3x 20 mins in PBST at room temperature, then incubated for 2 hours at room temperature with Alexa-Fluor secondary antibodies (ThermoFisher) (all 1:1500 in PBST + 5% BSA). Finally, embryos were washed 3x 20 mins in PBST at room temperature before being mounted in Vectashield containing DAPI (VectorLabs).

Oocytes were dissected from 2-day-old females. For Staufen and Gurken detection, 10 to 15 ovaries were fixed with PBS buffer containing 4% paraformaldehyde and 0.1% Triton X-100, washed three times for 5 mins in PBST and blocked in PBST containing 1% BSA. Incubation with the primary antibodies (anti-Staufen, Santa Cruz; anti-Gurken 1D12, DSHB) was performed overnight at room temperature or 4°C for Staufen and Gurken labelling, respectively, in PBT (PBS containing 0.1% BSA and 0.1% Tween 20). The ovaries were then briefly washed three times and three times for 30 min each in PBT and incubated for 2 hours at room temperature in Alexa-conjugated secondary antibodies. The ovaries were then washed 3x for 15 min each time in PBST, dissected, and mounted in Citifluor (Electron Microscopy Science).

### Phase contrast imaging of round spermatids

For analysis of round spermatids under phase contrast, testes were dissected in PBS, transferred to a 50μl droplet of PBS on a slide, cut open midway along the testes and, under observation, gently squashed under a coverslip using blotting paper.

### mRNA preparation and injection

pRNA vectors containing the appropriate cDNA were generated using Gateway cloning of PCR amplified cDNA and either a pRNA-GFP or a pRNA-mKATE backbone. pRNA vectors were linearised with AscI, precipitated using EDTA, sodium acetate, and ethanol, then resuspended in RNase-free water. mRNA was generated from these pRNA vectors *in vitro* using a T3 mMESSAGE mMACHINE kit (ThermoFisher) and was then purified using an RNeasy MinElute Cleanup kit (Qiagen). Freshly-laid unfertilised eggs were collected from apple juice plates within ∼1 hour of laying and were dechorionated on double-sided sticky tape. Eggs were lined up on heptane glue to keep them in place during injections and imaging. Embryos were dried at 25°C for ∼5 mins and covered with immersion oil (Voltalef). mRNA was manually injected using a syringe into eggs using needles made from borosilicate glass capillary tubes, at concentrations ranging from ∼2-4 μg/μl. Eggs were left for 1.5-2 hours before imaging to allow for translation of the mRNA. Control eggs were injected with RNAase-free water.

### Microscopy

Confocal imaging of fixed embryo (and the movies of scaffolds organising microtubules) was carried out on an Olympus FV3000 scanning inverted confocal system run by FV-OSR software using a 60X 1.4NA silicone immersion lens (UPLSAPO60xSilicone) or x30 0.95NA silicone immersion lens (UPLSAPO30xSilicone). Confocal imaging of scaffolds recruiting γ-TuRC proteins or organising microtubules and of testes samples was carried out on a Zeiss Axio Observer.Z1 inverted CSU-X1 Yokogowa spinning disk system with 2 ORCA Fusion camera (Hamamatsu) run by Zeiss Zen2 acquisition software using a 60X 1.4NA oil immersion lens (Zeiss). Confocal imaging of oocytes was carried out on a Zeiss LSM700 confocal microscope. Phase contrast microscopy of round spermatids was performed on a Leica DM IL LED inverted microscope controlled by μManager software and coupled to a RetigaR1 monochrome camera (QImaging) using a 40X 0.55NA air objective (Leica).

### Fertility tests

We tested fertility rates of males and females bred at 25°C, comparing pUbq-Cnn-C^T^ males or females to pUbq-Cnn-C males or females. We quantified the hatching rate of embryos that were generated when pUbq-Cnn-C or pUbq-Cnn-C^T^ males or females were crossed to *w^1118^* “wild-type” flies. Cages that were sealed with apple juice agar plates with a spot of dried yeast paste were set up at 25°C containing ∼50 newly-hatched test flies (e.g. pUbq-Cnn-C/ -C^T^) and ∼50 newly-hatched wild-type males or virgin females. The apple juice agar plates were exchanged with fresh plates 2-4 times a day, and the removed plates were kept at 25°C for at least 25 hours before the proportion of hatched eggs was calculated.

### Tissue Culture, Transfection, and IPs from HEK cells

HEK293T cells were grown in high glucose GlutaMAX Dulbecco’s modified Eagle medium (DMEM) supplemented with 10% heat inactivated foetal bovine serum and were incubated at 37°C and 5% CO_2_. Cells were mycoplasma free (LookOut Mycoplasma PCR detection kit, Sigma). Cells were passaged with 0.05% trypsin-EDTA every 2-3 days. 7x10^6^ cells were seeded and grown for 24 hours before transfection. Cells were transfected with 1.45 μg DNA using lipofectamine 2000 transfection reagent (ThermoFisher) for 4 hours in OptiMEM reduced serum medium. A control flask was treated with lipofectamine 2000 in the absence of any DNA but was otherwise processed identically. The medium was replaced with DMEM, and cells were allowed to grow for a further 16 hours before harvesting for immunoprecipitation.

Transfected cells were washed twice in PBS and lysed in buffer (50 mM HEPES, pH7.5, 150 mM NaCl, 1mM MgCL_2_, 1mM EGTA, 0.5% IGEPAL and protease inhibitors), and rotated for 90 minutes at 4°C. Cells were harvested at 15,000 rpm, 10 mins, 4°C. The supernatant was mixed with 30 μl GFP-Trap_MA beads (Chromotek) and rotated overnight at 4°C. Beads were washed 3 times in ice cold PBST, then resuspended in 50 μl 2x Laemmli sample buffer and boiled for 10 minutes at 95°C. Western blots were run as described above, using anti-GFP (1:250) (mouse, Roche) and anti-gamma-Tubulin (1:250) (rabbit, T5192 Sigma) primary antibodies.

### Image and statistical analysis

All images were processed using Fiji (ImageJ). *Quantifying and comparing the intensity of Cnn and γ-TuRC components at Cnn scaffolds*: Maximum intensity Z-plane projections were made and a threshold mask was generated using the Cnn channel. Sum fluorescence intensities for the Cnn and γ-TuRC protein channels were calculated. Overall mean cytosolic background intensity measurements for each channel were used to “background correct” the sum intensities for each scaffold. The scaffold intensities within each egg were plotted in Prism and a weighted linear regression analysis (based on ensuring an even distribution of residuals across the X axis) was performed. The gradient of the weighted regression line represented the S value for a given egg. The distribution of the S values per condition were lognormally distributed and so, to compare mean S values, the log_10_ of each individual S value was first calculated before performing an ANOVA analysis. This was to ensure the data being compared was normally distributed. Nevertheless, the unadjusted S values were plotted. Note that the fluorescence values and S values are in arbitrary units and cannot be directly compared between the γ-tubulin-mCherry and Grip75-sfGFP analysis. *Blind analysis of Cnn scaffolds organising microtubules*: Images were blinded ensuring each image had the same contrast settings. Images were then selected on scaffold size, with eggs containing small or very large scaffolds removed. The remaining images were then scored by eye as being of eggs that contained scaffolds with either no asters, weak or strong asters, or scaffolds where the Jupiter-mCherry signal did not extend beyond the GFP-Cnn signal (overlay). *Blind analysis of pUbq-Cnn-C or pUbq-Cnn-C^T^ embryos*: Images were blinded ensuring each image had the same contrast settings. Embryos were then scored by eye as being either normal or having moderate or severe defects. Embryos were scored as normal even when one or two mitotic figures had defects, because this is quite common in syncytial embryos and does not prevent development. Embryos were scored as having moderate defects when an unusually high proportion of mitotic figures had defects, or where the overall organisation of the spindles was moderately abnormal. Embryos were scored as having severe defects when there was either massive disorder with individual mitotic figures or overall organisation, or both. *To quantify western blot bands*: the sum intensities of bands were background corrected using mean “background” values at positions on the gel with no apparent signal. To reduce variation, the band intensities were taken using the freehand tool to draw closely around the perimeter of the band. For the Co-IP experiments, the intensities of the γ-tubulin IP bands were normalised to the intensity of the γ-tubulin band in the MBP-Cnn-T-N IP within each experiment. GraphPad Prism 7 or 8 was used for all statistical analysis and graph production.

### Bioinformatics

Protein alignments were produced using JalView. Secondary structure predictions were performed using JPred 4.

## Acknowledgements

This research was supported by a Wellcome Trust and Royal Society Sir Henry Dale fellowship awarded to PTC [105653/Z/14/Z], by IdEx Université de Paris ANR-18-IDEX-0001, by a Glover Fund research fellowship (Clare College/Dept. of Biochemistry, University of Cambridge) awarded to CAT, by an Association pour la Recherche sur le Cancer grant (PJA 20181208148) awarded to AG, and by the CNRS. We thank Jordan Raff for sharing Cnn antibodies and plasmids, Berthold Hedwig and Steve Rogers for help with needle pulling, Jens Lüders for sharing the pCMV-GFP vector, and Matt Castle for guidance on statistical analysis of Cnn scaffolds. We thank other members of the Conduit lab for their invaluable input and critical reading of the manuscript. The work benefited from the Imaging Facility, Department of Zoology, University of Cambridge, supported by Matt Wayland and a Sir Isaac Newton Trust Research Grant (18.07ii(c)), from the ImagoSeine at the IJM, Paris, and from use of the Cambridge Centre for Proteomics Core Facility. For the purpose of Open Access, the author has applied a CC BY public copyright license to any Author Accepted Manuscript version arising from this submission.

## Author Contributions

PTC and CAT designed the study and wrote the manuscript. CAT carried out cloning for all experiments and performed the *in vitro* recruitment assays, fertility tests, and round spermatid analysis. CT and PTC performed the mRNA injection experiments and analysed the scaffold data. PTC carried out the embryo analysis. AE helped establish the mRNA assay. AG performed the oocyte analysis and analysed the data. FB and AG generated the γ-tubulin-mCherry fly line via InDroso. MDR prepared bacterial cultures and assisted with protein purification.

## Supplementary Figure Legends

**Figure S1.**
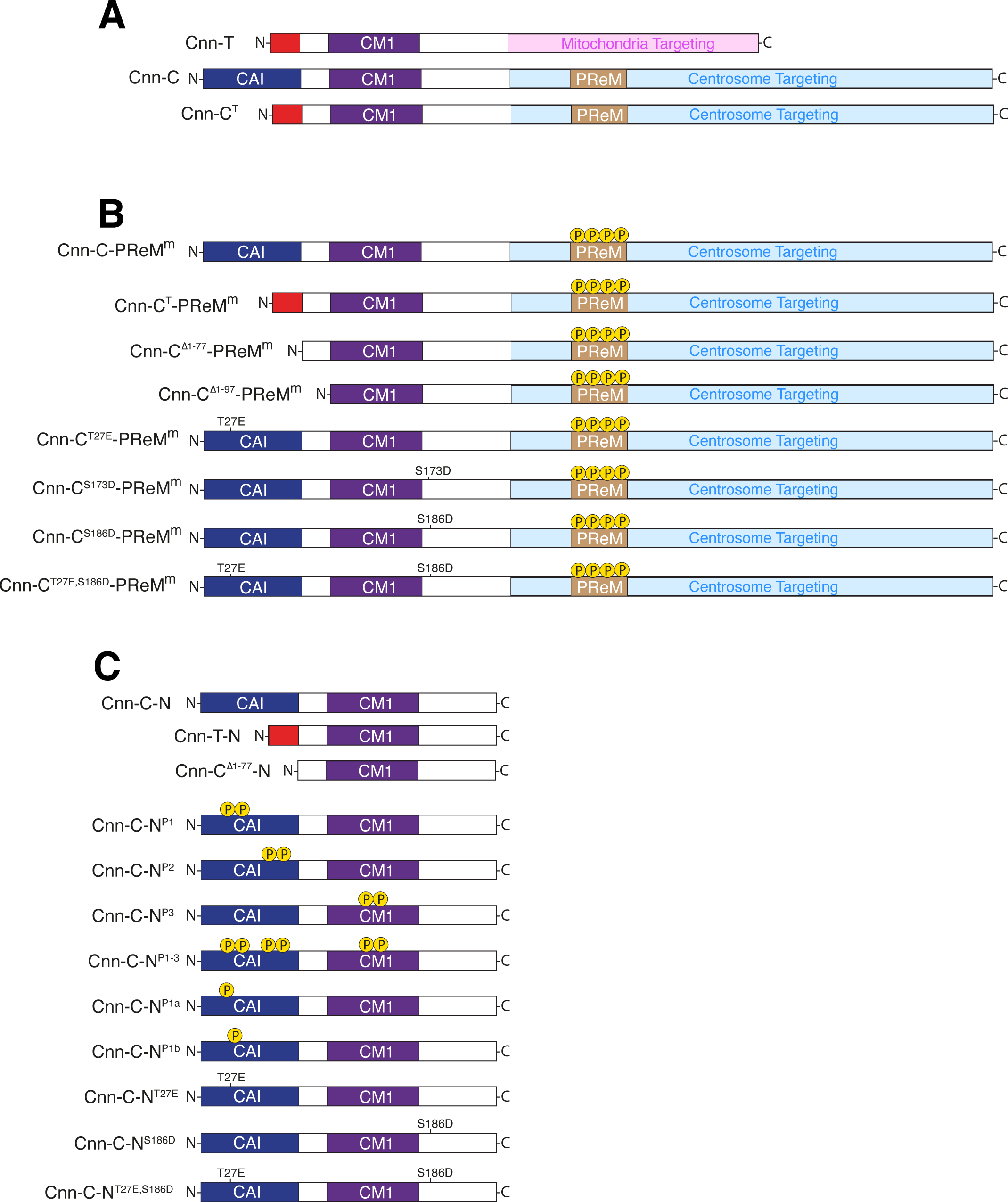
Diagrams of different Cnn constructs (omitting the tags) used in this study. **(A)** Diagram showing full-length Cnn constructs without modifications to the PReM domain. Cnn-T is the testes-specific isoform in *Drosophila*. Cnn-C is the major centrosomal isoform in *Drosophila*. Cnn-C^T^ represents an artificial form of Cnn-C in which the N-terminal region of Cnn-C (dark blue) has been replaced with the N-terminal region of Cnn-T (red). **(B)** Diagram showing Cnn constructs used in the scaffold assay, where Cnn-C contains phospho-mimetic mutations in the PReM domain to drive scaffold formation *in vivo*. **(C)** Diagram showing bacterially-purified N-terminal fragments of different Cnn types used in co-IP experiments.

**Figure S2.**
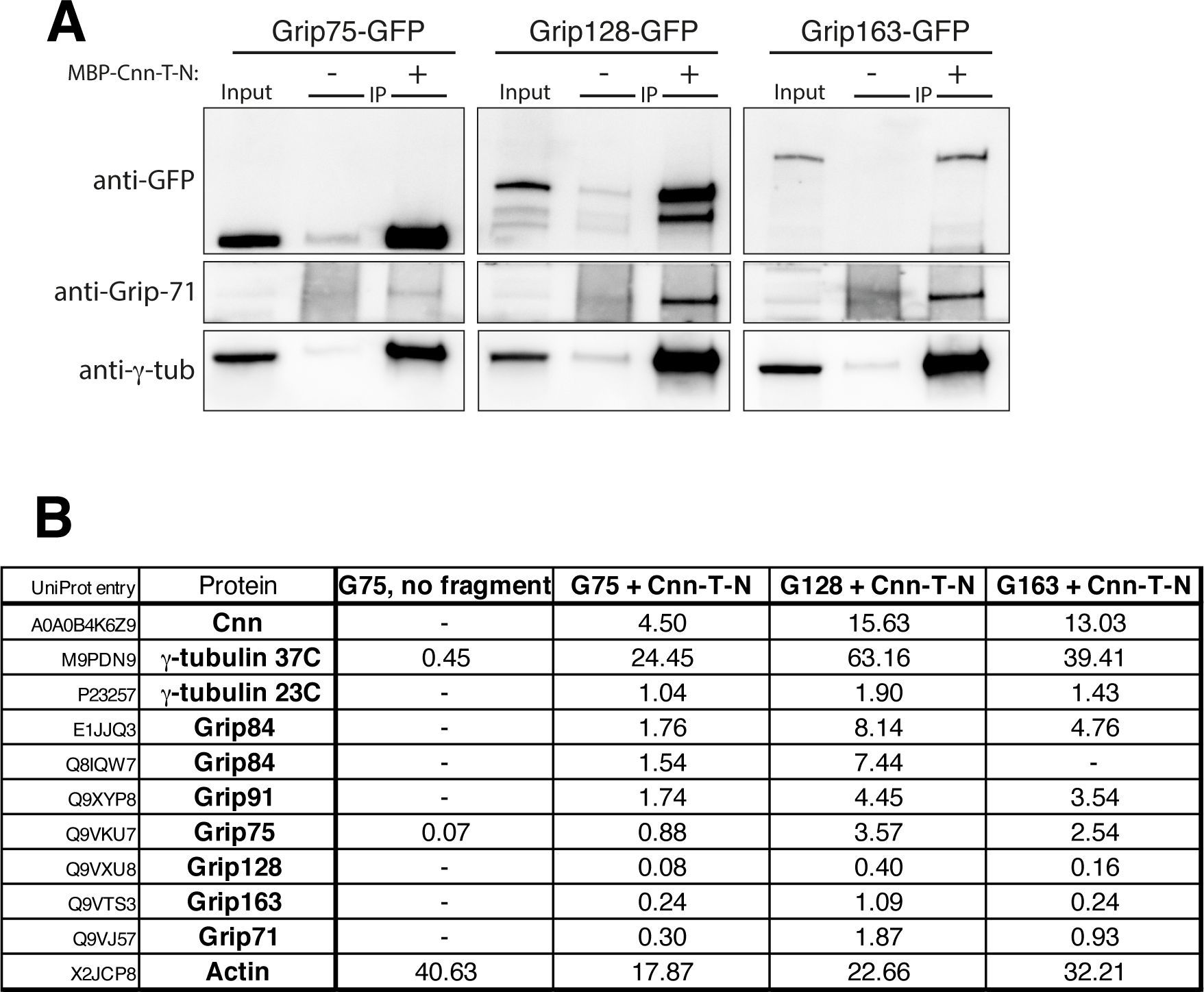
Bacterially-purified MBP-Cnn-T-N fragments immunoprecipitate γ-Tubulin Ring Complexes. **(A)** Western blot showing results of anti-MBP immunoprecipitation from embryo extracts expressing GFP-tagged Grip proteins (homologues of GCP4,5,6), either supplemented (+) or not supplemented (-) with MBP-Cnn-T-N, as indicated. Blots were probed with anti-GFP, anti-Grip71 and anti-γ-tubulin antibodies as indicated. When using MBP-Cnn-T-N, γ-tubulin and Grip71, as well as Grip75, 128, or 163, are co-immunoprecipitated. **(B)** Mass spectrometry results from IPs with MBP-Cnn-T-N showing the presence of various γ-TuRC components. Note that Mzt1 is not expressed within embryos. Results of a control experiment on Grip75-GFP embryo extract not supplemented with any MBP-Cnn-T-N fragment are also shown. Numbers indicate emPAI scores as a proxy for protein abundance. Grip84 (A) and Grip84 (E) represent two different isoforms of Grip84 (promoters 1 and 2 respectively).

**Figure S3.**
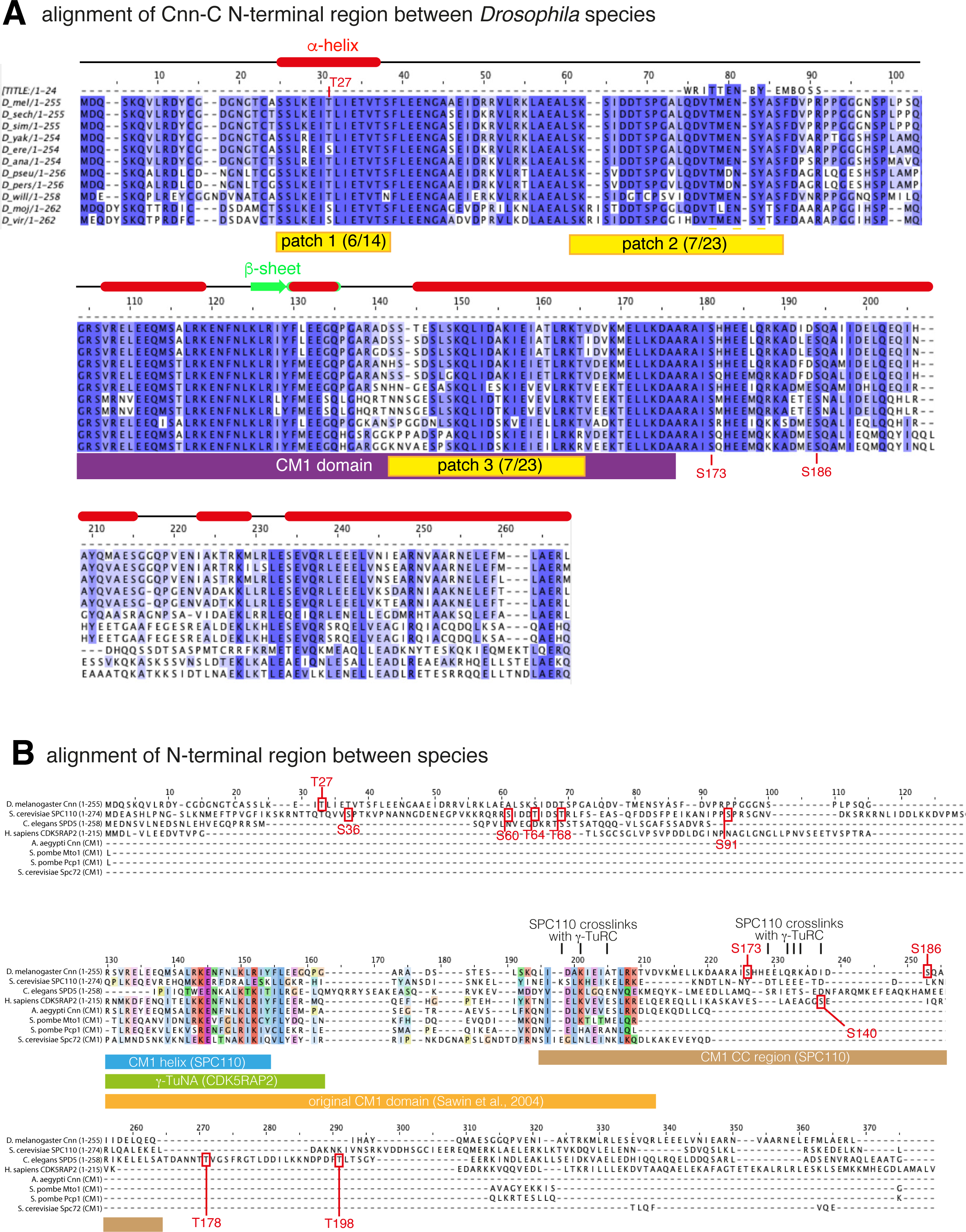
Protein alignments of N-terminal regions of CM1 domain proteins. **(A)** An alignment of Cnn-C homologues from different *Drosophila* species. The alignment was carried out in JalView keeping *D. melanogaster* at the top with the closest related species in order below. Only the N-terminal regions of the proteins were used in the alignment (∼1-255aa). Potential phosphorylation patches are highlighted in yellow, with the proportion of S/T residues present in the *Drosophila melanogaster* sequence indicated in brackets. The CM1 domain is highlighted in purple. Red boxes and green arrows indicate *α*-helices and *β*-sheets based on predictions from JPred. (**B**) An alignment of the N-terminal region of Cnn-C with the equivalent N-terminal regions of its homologues in non-*Drosophila* species. Phosphorylation sites that promote binding to γ-TuRCs identified either in this study (*Drosophila* Cnn-C, T^27^ and S^186^) or other studies (*S. cerevisiae* S^36^, S^60^, T^64^, T^68^ and S^91^,*C. elegans* SPD-5, T^178^, T^198^; human CDK5RAP2, S^140^) are indicated. Note that only the originally identified CM1 domain sequence (yellow) is conserved between homologues. The position of the “CM1 helix” (blue) and “CM1 coiled coil (CC) region” (brown) recently identified in SPC110 are indicated, as is the γ-TuNA sequence from human CDK5RAP2 (green).

**Figure S4.**
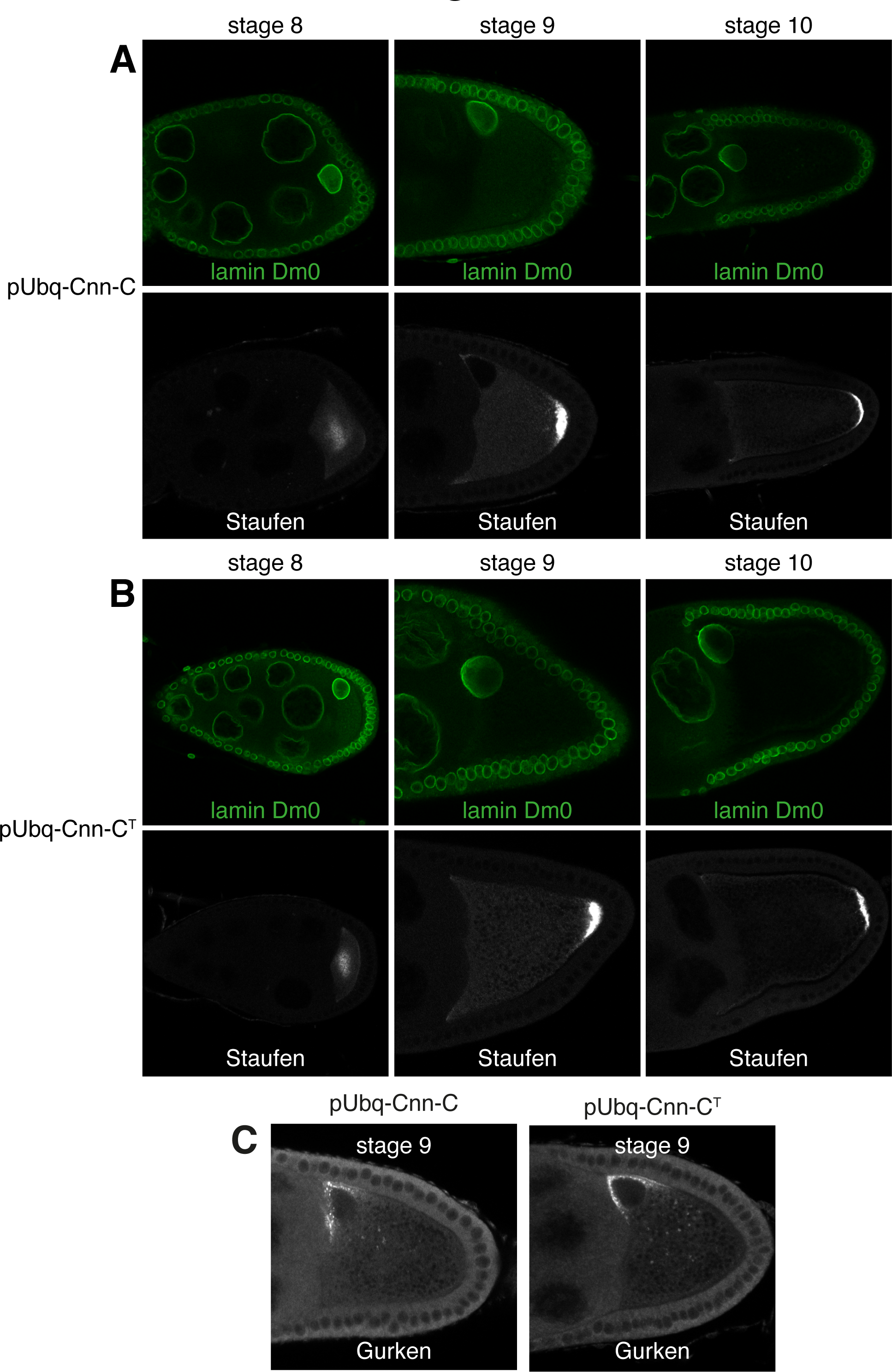
Polarity is established normally in pUbq-Cnn-C^T^ oocytes. **(A-B)** Fluorescence images show localisation of Staufen protein in oocytes expressing pUbq-Cnn-C (A) or pUbq-Cnn-C^T^ (B) at stages 8, 9 and 10, as indicated. Staufen localised in the centre of the oocyte at stage 8 and then at the posterior in stage 9 and 10 in all pUbq-Cnn-C (n=35, stage 8; n=35, stage 9; n=30, stage 10) and all pUbq-Cnn-C^T^ (n=40, stage 8; n=50, stage 9; n=40, stage 10) oocytes that were imaged. **(C)** Fluorescence images show localisation of Gurken protein in oocytes expressing pUbq-Cnn-C or pUbq-Cnn-C^T^ at stage 9. Gurken protein was positioned close to the nucleus in the dorsal corner in all pUbq-Cnn-C (n=30) and all pUbq-Cnn-C^T^ (n=35) stage 9 oocytes. Gurken mis-positioning or its absence results in abnormal dorsal appendages that protrude from the surface of the egg, but the dorsal appendages were normal on all pUbq-Cnn-C (n=724) and all pUbq-Cnn-C^T^ (n=488) eggs.

**Figure S5.**
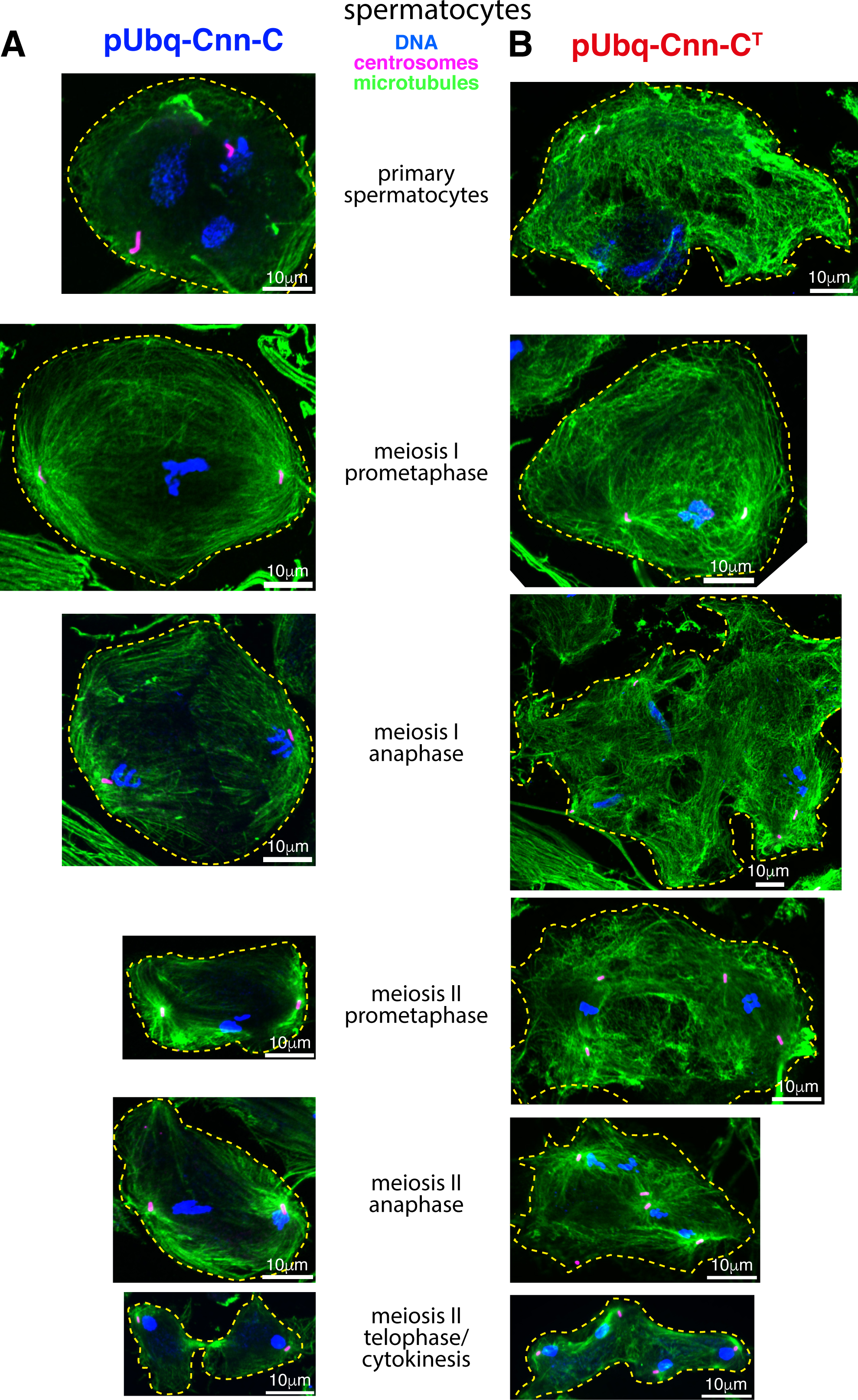
Major spermatocyte defects are observed within testes from pUbq-Cnn-C^T^ flies. (**A,B**) Fluorescence images showing spermatocytes at different developmental stages (as indicated) from flies expressing either pUbq-Cnn-C (A) or pUbq-Cnn-C^T^ (B) fixed and stained for microtubules (green, *α*-tubulin), centrosomes (pink, Asterless), and DNA (blue). A high density of cytosolic microtubules, as well as cytokinesis and karyokinesis defects, are clearly observed in cells from pUbq-Cnn-C^T^ testes.

## Supplementary Videos

Video 1

**Cnn-T scaffolds organise microtubule asters and can be mobile.** Movie showing Cnn-T scaffolds (green) organising microtubule asters (marked with Jupiter-mCherry (magenta)). A mobile scaffold (lower left) with an asymmetric microtubule aster can be seen moving through the cytosol.

Video 2

**Transient spindle-like structures can form between Cnn scaffolds.** Movie showing the formation and disappearance of a transient spindle-like structure between adjacent Cnn-T scaffolds (green). Microtubules are marked with Jupiter-mCherry (magenta).

Video 3

**Spindle-like structures organised by Cnn scaffolds can form in synchrony.** Movie showing the synchronous formation and disappearance of a multi-polar spindle-like array of microtubules that is subsequently organised by a nearby group of coalescing Cnn scaffolds (green). Microtubules are marked with Jupiter-mCherry (magenta)

Video 4

**Microtubules are robustly anchored to Cnn scaffolds.** Movie showing rare giant Cnn-T scaffolds (green). One scaffold can be seen rotating and dragging the microtubules, indicating that the microtubules are robustly attached to the scaffold, presumably via γ-TuRCs. Microtubules are marked with Jupiter-mCherry (magenta).

Video 5

**Expression of GFP-Cnn-T-N leads to the formation of dynamic microtubules within the cytosol of unfertilised eggs.** Video shows the effect of injecting mRNA encoding GFP-Cnn-T-N into unfertilised eggs expressing Jupiter-mCherry (marker of microtubules). Left panel shows the GFP channel (green), centre panel shows the RFP channel (magenta), right panel shows a merge.

## Notes

### Competing Interest Statement

The authors have declared no competing interest.

### Summary of Updates

Data has been added (new Figure 4) on how phospho-mimetic mutations at T27 and S186 within Cnn promote recruitment of g-TuRCs to Cnn scaffolds, and on how these mutations dramatically increase binding to g-TuRCs in co-IP assays. Data has been added (new Figure 8) on how the region downstream of the CM1 domain in human CDK5RAP2 inhibits binding to g-TuRCs.

